# Learning the mutational landscape of the cancer genome

**DOI:** 10.1101/2021.08.03.454669

**Authors:** Maxwell A. Sherman, Adam Yaari, Oliver Priebe, Felix Dietlein, Po-Ru Loh, Bonnie Berger

## Abstract

An ongoing challenge to better understand and treat cancer is to distinguish neutral mutations, which do not affect tumor fitness, from those that provide a proliferative advantage. However, the variability of mutation rates has limited our ability to model patterns of neutral mutations and therefore identify cancer driver mutations. Here, we predict cancer-specific mutation rates genome-wide by leveraging deep neural networks to learn mutation rates within kilobase-scale regions and then refining these estimates to test for evidence of selection on combinations of mutations by comparing observed to expected mutation counts. We mapped mutation rates for 37 cancer types and used these maps to identify new putative drivers in understudied regions of the genome including cryptic alternative-splice sites, 5’ untranslated regions and infrequently mutated genes. These results, available for exploration via web interface, indicate the potential for high-resolution neutral mutation models to empower further driver discovery as cancer sequencing cohorts grow.

## Introduction

Neutral (passenger) mutations that do not provide a proliferative advantage to a cell dominate the mutational landscape of tumors^1,2^. Only a relatively small fraction of mutations that target specific genomic elements (e.g., coding sequence, promoters, enhancers, and lncRNAs) are under positive selection^3–5^ due to their ability to promote cell growth, resist cell death, or enable tissue invasion^6^. Efforts to distinguish positively-selected driver mutations, genes, and noncoding elements from neutral passengers have informed our understanding of somatic mutation processes and cancer disease mechanisms^4,5,7–12^. Nonetheless, catalogs of cancer driver elements remain incomplete – particularly in the noncoding genome^13^ – hindering precision oncology^4,8,14,15^.

Positive selection on driver elements results in more mutations than expected compared to the rates at which neutral mutations occur^16,17^. Robust identification of such signals thus requires an accurate computational model of the neutral mutation rate within a genomic element of interest, a task made challenging by the highly variable, tissue-specific patterns of neutral mutations across the cancer genome^18,19^. Currently, computational methods for driver detection model the neutral mutation rate in specifically defined regions of the genome^17,20–22^, which has been particularly successful in the coding genome where synonymous mutations can serve as a proxy for neutral mutations^3,4,23,24^. However, cancer driver mutations may exist anywhere in the genome and in patterns that may not be definable as standard genomic intervals; noncoding driver mutations can be spread widely across a driver gene’s regulatory elements and may not be identifiable by examining any single element in isolation^25–27^. Moreover, whole-genome sequencing of tumor genomes can assay passenger and driver mutations genome-wide^9,28,29^. These considerations motivate the development of high-resolution neutral mutation rate models that span the entire cancer genome.

Here, we introduce a probabilistic deep-learning framework that estimates neutral somatic mutation rates genome-wide through a two-step process. First, a deep neural network predicts kilobase-scale somatic mutation rates for a cancer of interest using high-resolution epigenetic organization of healthy tissues (a well-known predictor of tumor mutation rates at the megabase scale^18,30^). Then, by leveraging the nucleotide context biases of cancer-specific mutational processes, these kilobase-scale predictions are refined via a hierarchical probabilistic model to estimate mutation rates within finer-scale genomic elements with state-of-the-art accuracy. This approach enables *any combination of candidate mutations* to be efficiently evaluated for evidence of selection by testing for a burden of observed mutations using the resources of a personal computer. We mapped cancer-specific neutral somatic mutation rates using the Pan-Cancer Analysis of Whole Genomes (PCAWG) dataset^9^, and we used these mutation models to identify new candidate cancer drivers by analyzing publicly available whole-genome, whole-exome, and targeted-sequencing cancer datasets. Our mutation maps of 37 different cancers are publicly available both as an interactive genome-browser and as a standalone software tool for quantifying excess somatic mutations anywhere in the genome in a dataset of interest.

## Results

### Modeling somatic mutation rates from epigenomic features and nucleotide context

Neutral somatic mutation rates vary dramatically over different scales of the cancer genome^18,19,23^. Higher-order genomic properties such as replication timing and chromatin accessibility reflect mutation rate variability at the megabase scale due in large part to differential DNA repair in late versus early replicating regions and euchromatic versus heterochromatic regions^17^. However, less is understood about the relationship between local epigenetic structures and kilobase-scale variation in mutation patterns^19,31,32^. To explore the extent to which finer-scale epigenomic profiles might be informative of genome-wide mutational landscapes (dominated by neutral mutations) observed in cancer cohorts, we designed a deep-learning method to predict somatic mutation rates and quantify prediction uncertainty in kilobase-scale regions of cancer genomes using 100bp-resolution chromatin patterns in healthy tissue from the Roadmap Epigenomics^33^ and ENCODE^34^ datasets (**Fig. 1a**, **fig. S1**, **table S1**).

**Figure 1:**
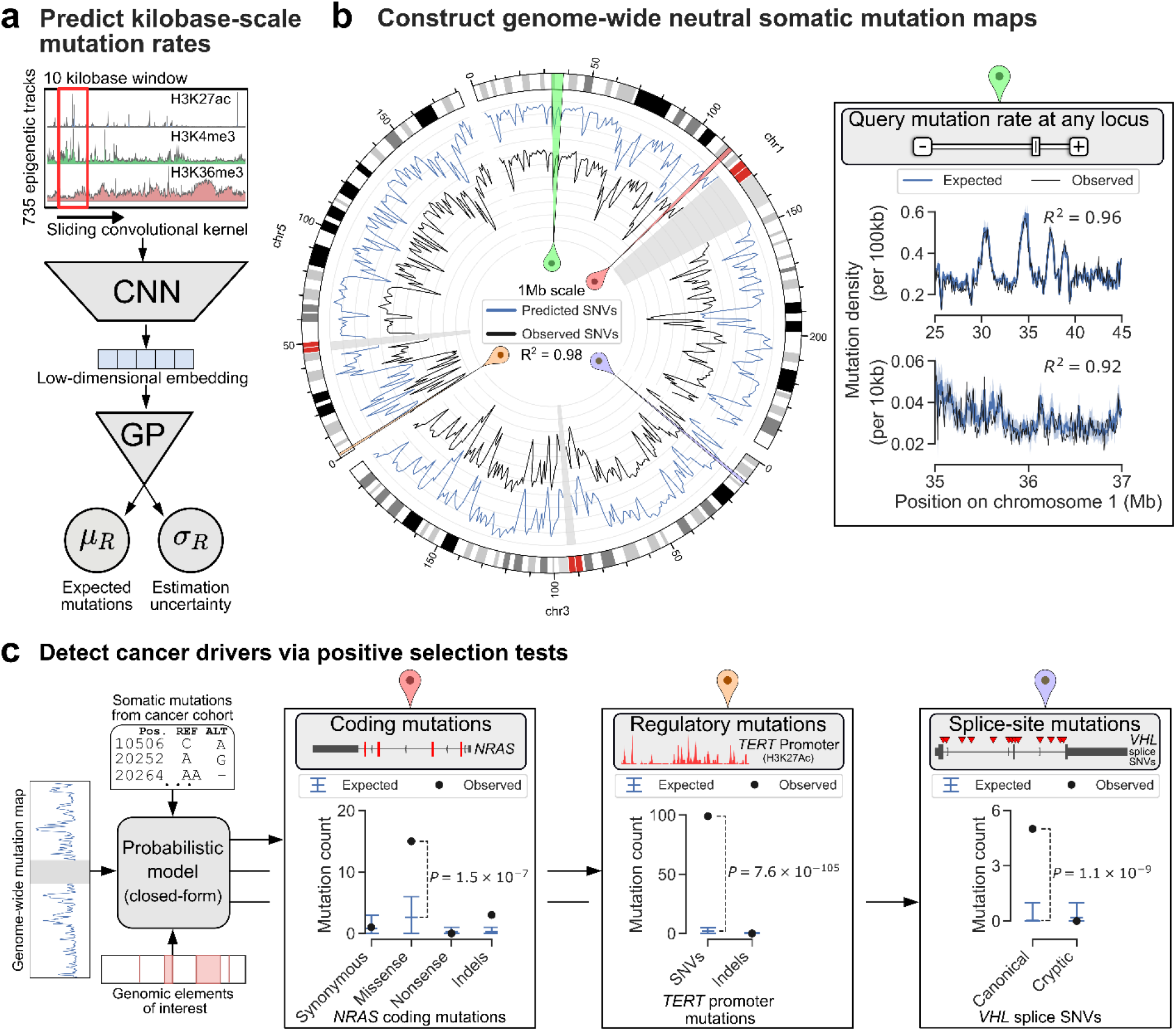
Modeling the genome-wide neutral somatic mutation rate and identifying cancer driver elements. **a,** Deep-learning scheme used to predict the expected number of somatic mutations along with the prediction uncertainty using high-resolution epigenetic profiles of healthy tissue from the Roadmap Epigenomics consortium and ENCODE. CNN: convolutional neural network; GP: Gaussian process. **b,** Genome-wide neutral somatic SNV map and the observed density of SNVs in 1Mb windows from the PCAWG pan-cancer cohort (n=2,279 samples). For clarity, only chromosomes 1, 3, and 5 are shown. Highlighted are regions of interest demonstrating the ability to estimate the neutral mutation distribution in any genomic locus. Inset shows a region on chromosome 1 modeled at 100kb resolution and 10kb resolution. The reported R-squared statistic between observed and expected SNV counts is calculated genome-wide for the associated resolution. **c,** The genome-wide neutral mutation map for a cancer of interest is integrated with (i) a dataset of somatic mutations from a cancer cohort and (ii) user-defined genomic elements, using a probabilistic model to estimate a closed-form distribution over the expected number of neutral mutations in each element. Positive selection is then tested by calculating the likelihood of the observed mutation under the null hypothesis of this neutral mutation distribution. Genomic elements to test (indicated in red) can be any sets of contiguous or noncontiguous regions (e.g., coding sequence and regulatory regions), or sets of mutations occurring at specific base pairs.

We benchmarked how well our method predicted local mutation rates on 37 cancer types from the PCAWG dataset (**table S2**) in five-fold cross-validation analyses that strictly partitioned the genome into non-overlapping train and test sets (predicting mutation rates in each one-fifth of the genome using a model trained on observed mutations in the remaining four-fifths; **Methods**). Our method successfully predicted a median of 71.2% (range: 17.0-90.0%) of variance in observed single nucleotide variant (SNV) rates in 10kb regions and a median of 94.6% (range: 73.1-98.0%) of variance in 1Mb regions (**Fig. 1b**, **table S3**) across 16 cancer types for which benchmarking power was sufficient (>1 million mutations and excluding lymphomas; **Methods**). Including mutation counts in flanking regions as additional input to the model provided a small but consistent increase in accuracy (**table S3**) (median variance explained in 10kb regions: 77.3%; range 22.7-92.3%), suggesting that mutational processes not reflected in local epigenetic structures of healthy tissue play a subtle but meaningful role in kilobase-scale mutation rate variability.

To gain insight into which specific epigenetic features the deep-learning model utilized to predict mutation patterns, we leveraged an approach that highlights input features important to the model’s accuracy (feature maps, **Methods**). The deep-learning network consistently focused on localized features (avg. size 1526 bp; 95%CI: 1512-1540 bp) corresponding to transcription start sites, regions of active transcription, enhancers, repressive regulatory states, and heterochromatin to make predictions within kilobase-scale regions (**fig. S2-S5**). These results are consistent with known epigenetic correlates of mutation rates ^19^ and add to the growing evidence that deep-learning models can extract and encode biologically relevant signals from high-dimensional experimental data^35–37^.

We next sought to determine if the neutral mutation models could re-identify driver genomic elements in which mutations are known to be under positive selection in cancers, producing an excess of observed mutations relative to expectation (**Fig. 1c**). The successes of existing methods to test for mutational excess within predefined genomic intervals covering specific portions of the genome^5,17,20–22^ motivated us to build a method that could flexibly and efficiently test for excess across any set of candidate mutations genome-wide. We thus devised a method to approximately subdivide kilobase-scale neutral mutation rate predictions from the deep learning model into base-pair-level predictions. Specifically, sequence context is the major determinant of neutral mutation rates at base-pair resolution^7,19,38,39^; therefore, we partitioned kilobase-scale mutation rate maps into approximate base-level estimates according to observed trinucleotide mutational signatures, using the relative likelihoods of mutations based on sequence context to estimate how mutations should be distributed across positions within a kilobase-scale region (**Methods**).

This approach of regional rate estimation (based on epigenetics) followed by distribution refinement (based on mutational signatures) successfully modeled mutation rates in finer-scale elements, explaining as much or more variance in observed mutation rates as bespoke mutation rate models developed for specific genomic elements (e.g., tiled regions, coding sequence, enhancers, and noncoding RNAs) (**fig. S6**; **table S3-S6**). The accuracy of the mutation rate models enabled correspondingly powerful driver identification: in benchmarks testing downstream ability to identify evidence of positive selection (i.e., excess of mutations) on previously-identified driver mutations in various types of genomic elements, the approach matched or exceeded the performance of previous methods tailored toward specific classes of elements^4,20–23^ (e.g., coding sequences and promoters; **figs. S7-S9**, **table S7-S8**, **Note S1-S2**). Moreover, testing for mutational excess within 10^5^ genomic elements completed in two minutes on a single CPU core (**fig. S10**) due to the closed form we derived for the neutral mutation distribution estimated for combinations of candidate mutations (**Methods**).

To maximize accessibility of the mutation rate maps we generated, we have created a web-based genome-browser resource where expected and observed counts from 37 types of cancer from the PCAWG dataset can be visually browsed and compared (**Data availability**) (**fig. S11**).

### Quantifying evidence of selection on cryptic splice SNVs across cancers

We statistically analyzed the neutral mutation maps together with observed mutation counts in PCAWG to identify sets of mutations manifesting evidence of positive selection (mutational excess) in several understudied regions of the genome. We first considered mutations that may promote cryptic alternative splicing. Alternative-splicing is increasingly recognized as functionally relevant to cancer^40,41^ and recent studies have associated specific somatic mutations outside the canonical splice sites with alternative splicing events observed in expression data^42,43^. These associations motivated us to rigorously quantify the extent to which cryptic splice SNVs occur in excess of the neutral mutation rate and therefore may function as driver mutations under selection.

We thus applied our method to map mutation rates and test for mutational excess at cryptic splice sites, which may occur widely in both exons and introns of a gene^44^. We analyzed potential splice-altering SNVs predicted by SpliceAI^44^, which uses a deep-learning model together with *in-silico* mutagenesis to assess the splicing impact of all possible SNVs within transcribed gene bodies. We stratified predicted splice-altering SNVs into canonical and cryptic splice mutations and by predicted splicing impact (SpliceAI Δ score) (**Fig. 2a**) (**Methods**), and we calculated enrichments of observed SNV counts in PCAWG relative to expectation under our neutral model (**Methods**). In known tumor suppressor genes (TSGs), we observed cryptic splice SNVs significantly more often than expected under neutrality (**Fig. 2b**, **table S9**). The enrichment was driven largely by intronic cryptic splice SNVs (which comprise the majority of such mutations) and increased with predicted impact, with mutations in the highest-impact category (Δ score > 0.8) exhibiting 1.75-fold enrichment (95% CI: 1.31-2.22 fold; P=2.52×10^-5^), roughly half as large as for canonical splice SNVs (**Fig. 2b**). These results were robust to the inclusion or exclusion of high mutation burden samples (**fig. S12**) and consistent with an analysis that did not rely on the mutation maps (**fig. S13**). A control analysis of oncogenes and genes not in the Cancer Gene Census (CGC)^45^ did not find enrichments of predicted loss-of-function (pLoF) SNVs or cryptic splice SNVs (**fig. S14**, **table S9**).

**Figure 2:**
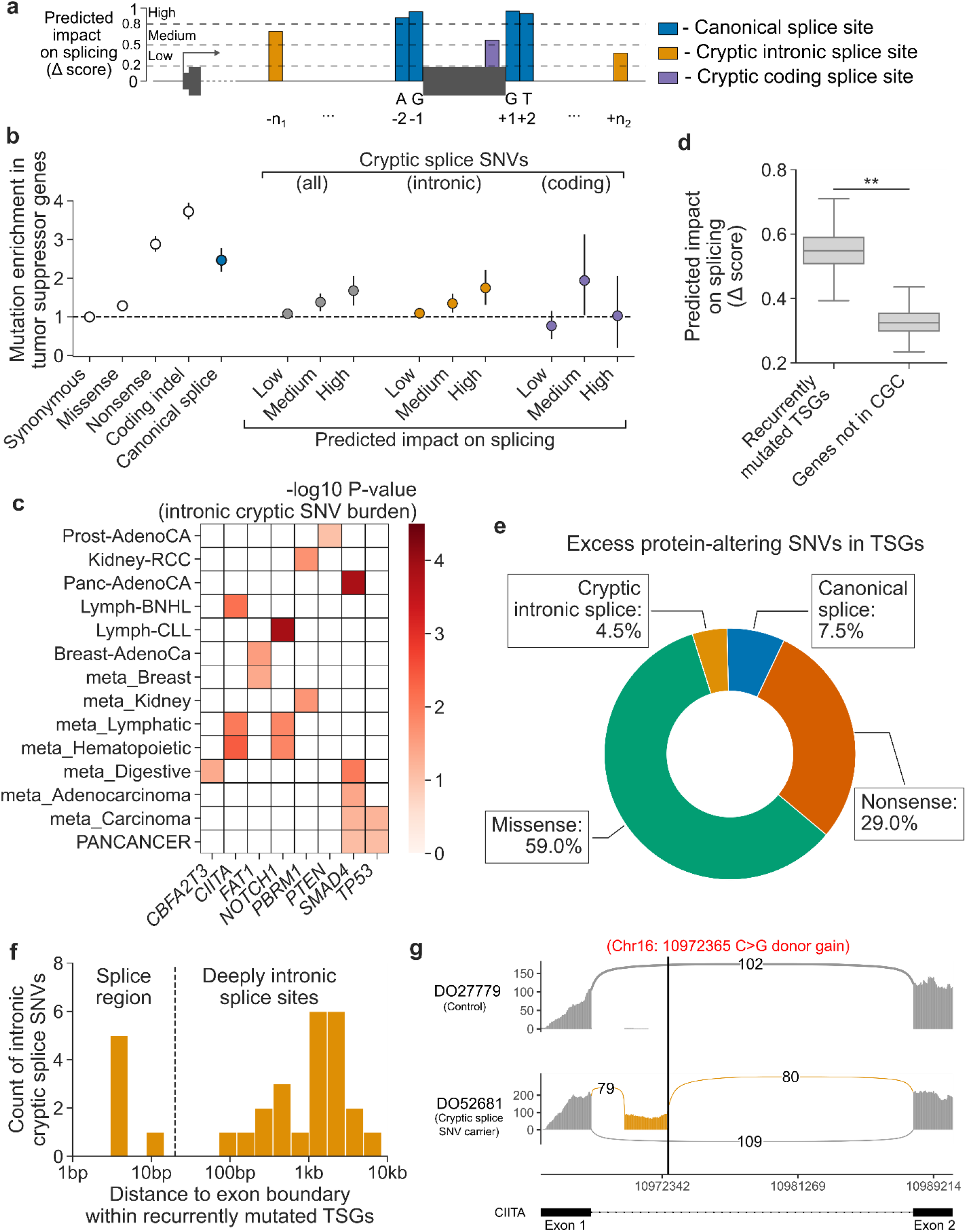
Evidence of positive selection on intronic cryptic splice SNVs in tumor suppressor genes. **a,** A schematic of the splice-altering SNVs considered in this analysis. Predicted impact on splicing is measured by the SpliceAI Δ score (higher score corresponds to higher predicted impact). We further stratified predicted splice-altering SNVs by predicted impact on splicing: low predicted impact (0.2<Δ<0.5)’ medium predicted impact (0.5<Δ<0.8) and high predicted impact (0.8<Δ<0.1). **b,** Enrichment (with 95% confidence interval) of observed mutations compared to expected neutral mutations in tumor suppressor genes stratified by variant type and predicted impact on splicing. **c,** Known tumor suppressor genes per cancer with a significant burden (FDR<0.1) of predicted intronic cryptic splice SNVs. **d,** Predicted splicing impact (SpliceAI Δ score) for intronic cryptic splice SNVs observed in recurrently mutated TSGs (**c**) compared to those observed in genes not in the Cancer Gene Census (CGC) (bootstrapped P<3×10^-4^). **e,** Proportion of excess SNVs in TSGs contributed by each protein-altering SNV category. **f,** Distribution of distance to nearest exon boundary for the intronic cryptic splice SNVs observed in recurrently mutated TSGs. **g,** Pileup of RNA-seq reads in a Lymph-BNHL carrier of a predicted, deeply intronic cryptic splice SNV (labeled in red) in *CIITA* and a control Lymph-BNHL sample, showing the inclusion of a cryptic exon (gold) in the cryptic splice SNV carrier. Arc labels indicate the number of RNA-seq reads that support each exon junction.

A search for individual TSG-cancer pairs contributing to this enrichment identified eight TSGs in 14 cohorts with a significant burden (FDR<0.1 for n=283 TSGs in 37 cancer types) of intronic cryptic splice SNVs (**Methods**) (**Fig. 2c**, **table S10**). The pattern of associations between specific TSGs and types of cancer was consistent with known tissue specificity of TSGs. In pan-cancer analyses, predicted intronic cryptic splice sites were recurrently mutated in *TP53* and *SMAD4,* which have been implicated in numerous cancers; in contrast, the hematopoietic-specific TSG *CIITA* was recurrently mutated in blood malignancies and the renal-specific TSG *PBRM1* was recurrently mutated in kidney malignancies. In further support of these associations, the intronic cryptic splice SNVs observed in these recurrently altered TSGs had significantly higher predicted impact on splicing than those observed in genes not in the CGC (**Fig. 2d**) (mean SpliceAI Δ score = 0.55 vs. 0.33; P < 3×10^-4^; **Methods**).

We next quantified the contribution of intronic cryptic splice SNVs to candidate TSG driver events across cancers. To do so, we estimated the excess of cryptic splice SNVs relative to the excess of predicted protein-altering SNVs (including coding and canonical splice mutations) across all TSGs in the CGC (**Fig. 2e**, **Table S11**) (**Methods**). Intronic cryptic splice SNVs accounted for 4.5% (95% CI: 1.3-7.4%) of excess SNVs, similar in magnitude to the 7.4% (5.6-9.7%) attributable to canonical splice SNVs. Missense SNVs accounted for 59.0% (53.7-63.7%) of excess SNVs and nonsense mutations accounted for 29.0% (25.6-32.9%). As expected, we did not observe an excess of synonymous mutations (1502 observed vs. 1503 expected; P=0.51). Excess SNV estimates were consistent with excess estimates from dNdScv when restricting the analysis to protein-coding and canonical splice SNVs (**fig. S15**).

The majority (79.3%) of intronic cryptic splice SNVs in recurrently altered TSGs fell outside annotated splice regions (i.e., >20bp from exon-intron boundaries) (**Fig. 2f**). To further asses the function of deeply intronic mutations, we used LeafCutter^46^ to search for cryptic splicing in RNA-seq data available for nine cryptic splice SNV carriers (**Methods**). Of the seven carriers with sufficient coverage, six had evidence of alternative splicing (**Fig. 2g**, **fig. S16**, **table S12**, **Note S3**). Overall, these results provide evidence that intronic cryptic splice SNVs are under positive selection in TSGs and likely act as driver events in an appreciable fraction of tumors (several percent) across multiple cancer types.

### Noncoding candidate cancer driver mutations in 5’ UTRs of TSGs

We next searched for potential driver mutations in promoter regions, another class of noncoding genomic elements in which mutations can have large effects on gene function^47–49^. We first applied our model only to indels, hypothesizing that they may have larger effect sizes in noncoding regulatory elements than SNVs because they can disrupt larger portions of binding motifs. Searching all promoters (n=19,251) for evidence of selection on indels in the PCAWG pan-cancer dataset (**Methods**) revealed one element with a Bonferroni-significant burden of indels: the promoter of *TP53* (7 observed vs. 0.54 expected; P = 9.4×10^-7^). While previous analysis of the PCAWG dataset (using restricted hypothesis testing to boost statistical power) also observed a significant (*Q*=0.044) recurrence of *TP53* promoter mutations ^5^, the strength of the indel enrichment we observed here motivated further exploration of this signal.

The observed mutations – all deletions – specifically affected exon 1 of the canonical 5’ UTR and disrupted sequence elements critical to transcription (e.g., transcription start site), translation (*WRAP53* binding sequence^50^ and internal ribosome entry site^51,52^), and splicing within the 5’ UTR (which spans multiple exons) (**Fig. 3a**). The deletions were significantly larger than most 5’ UTR indels in tumors (**Fig. 3b**) (median length = 17bp vs 1bp; P=7.4×10^-4^, Mann-Whitney U-test) and exhibited enrichment comparable to cryptic exonic splice SNVs in *TP53,* which are well-characterized cancer drivers^53^ (**Fig. 3c**). While the splice regions of many *TP53* exons were recurrently mutated, the pattern of mutations in the exon 1 splice donor region appeared to be unusual, with more than half of mutations (four of seven) not altering the canonical splice sites (**Fig. 3d**) (P=1.8×10^-3^, Fisher’s exact test). RNA-seq data revealed that carriers had 1-2 standard deviation decreases in *TP53* expression (**Fig. 3e**) (P=1.6×10^-4^, **Methods**). Evidence of alternative splicing was not present in transcripts sequenced from the splice region mutation carriers, suggesting that the 5’ UTR splice mutations resulted in nonsense-mediated decay, possibly due to open-reading frames within the retained intronic sequence. To further corroborate these results, we examined somatic mutations in 2,399 distinct samples from the Hartwig Medical Foundation^54^. These samples showed a similar mutational pattern, with three samples carrying >10bp deletions and four samples carrying SNVs in exon 1 of the *TP53* 5’ UTR and its donor splice region (**Fig. 3a**).

**Figure 1:**
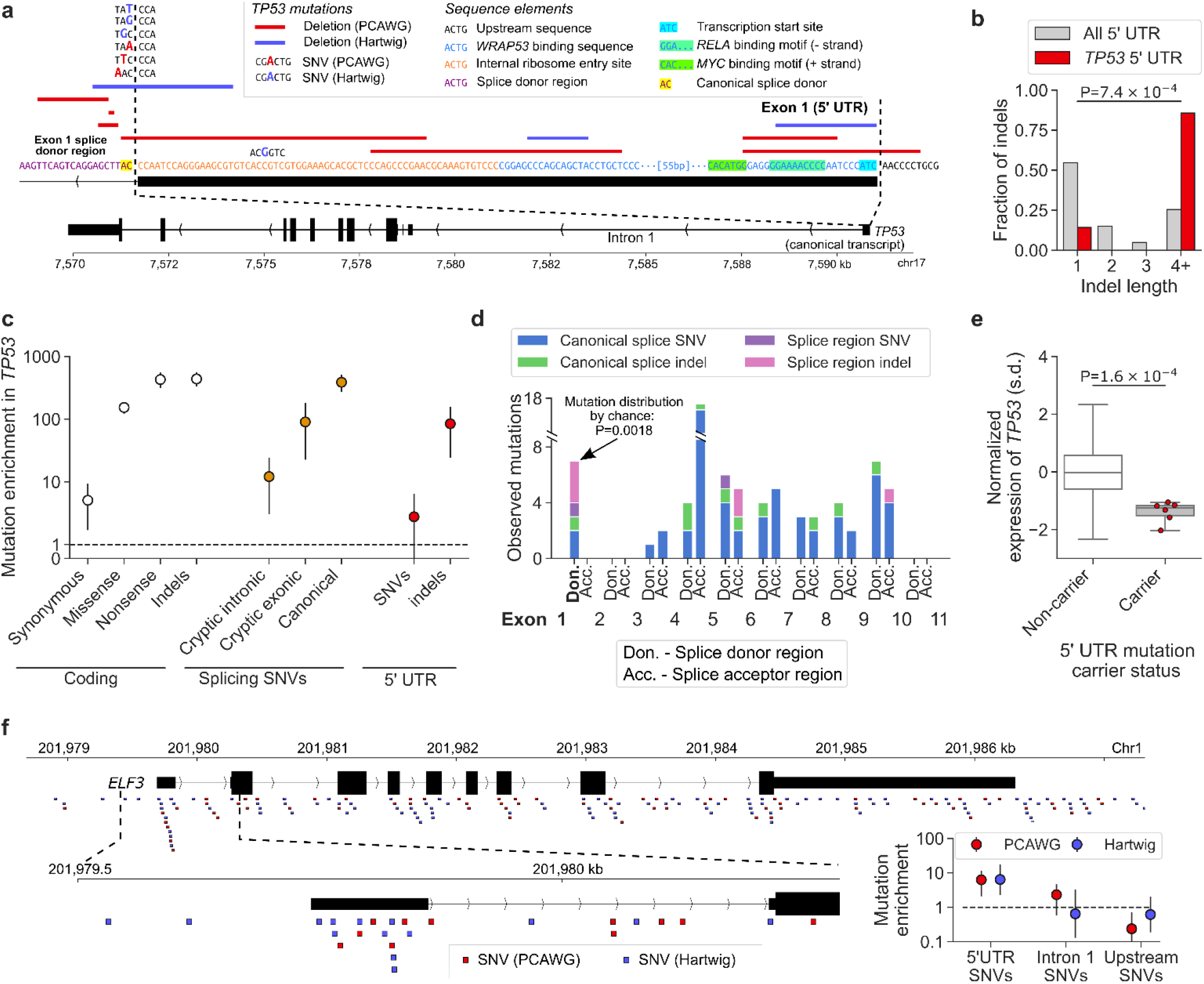
Enrichment of somatic mutations in the 5’ UTRs of *TP53* and *ENF3.* **a**, Mutations observed within exon 1 of the 5’ U TR of the canonical *TP53* transcript from the PCAWG and Hartwig Medical Foundation cohorts. The DNA sequence shown is from the GRC h37 reference genome (+ strand). Mutation types, relevant sequence and reg ulatory elem ents are indicated in the legend. **b,** Distribution of indel sizes obserned within 5’ UTRs of genes other than *TP53* and within tha *TP53* 5’ UTR. P-value comparing median indel lengths, Mann-Whitney U-test. c, Mutation enrichment relative to the neutral mutation rate (observed / expected neutral mutations) within *TP53* stratified by mutation type and location. Error bars, 95% CI. **d,** Distribution of mutations observed within the donor and acceptor splice regions (defined as the 20bp 3’ and 5’ of an exon, respectively) of the canonical *TP53* transcript. Canonical splice SNVs and indels are defined as mutations that alter the two base-pairs immediately adjacent to an exon boundary; splice region SNVs and indels are defined as mutations that intersect the splice region but do not disrupt the canonical splice sites. The donor splice region of exon 1 of the 5’ UTR (shown in **a**) is bolded. The P-value of observing the distribution of canonical and splice region mutations in the donor splice region of exon 1 5’ UTR compared to all other *TP53* splice regions was computed via a Fisher’s exact test. **e,** Expression of *TP53* on standard deviation (s.d.) scale in carriers of *TP53* 5’ UTR mutations (n=6) and non-carriers (n=1,205), adjusted for tumor type and copy number. P-value via Mann-Whitney U-test on adjusted and standardized expression values. **f,** SNVs overlapping *ELF3* in the PCAWG and Hartwig Medical Foundation cohorts. Insets: zoom-in of the *ELF3* 5’ UTR region and mutational enrichments within this region.

These results motivated a targeted search for mutational burden in 5’ UTRs and their splicing regions across 106 TSGs and 95 oncogenes with multi-exonic 5’ UTRs (**Methods**). The analysis revealed one additional element with a significant burden of SNVs or indels: the 5’ UTR of *ELF3* (**Fig. 3f**) (6 observed SNVs vs 0.96 expected; P=2.9×10^-4^). Samples from the Hartwig Medical Foundation displayed a similar enrichment of SNVs in the *ELF3* 5’ UTR (10 observed vs. 1.5 expected; P=3.8×10^-4^, **Methods**). In both sets of samples, the enrichment was concentrated within the canonical *ELF3* 5’ UTR; surrounding sequences (upstream promoter and intron 1) were not enriched for mutations (**Fig. 3f**). Interestingly, the 16 total mutations largely altered distinct base pairs within the 5’ UTR – although two positions mutated in PCAWG samples were also mutated in the Hartwig samples – suggesting that this 5’ UTR might be broadly sensitive to perturbation, possibly by prompting changes in promoter methylation that alter *ELF3* expression^55^. An alternative possibility could be an unmodeled local mutational process or technical artifact in this region^17^; however, a careful analysis did not find evidence for any such features that have explained other noncoding mutational hotspots^5^ (**Note S4**). Additional follow-up will be necessary to determine whether the mutational enrichment here represents positive selection or points to a new neutral mutational process.

### The shared landscape of common and rare driver genes

Clinical reports have revealed that tumors of one cancer type occasionally carry mutations frequently implicated as drivers in another cancer type^56,57^. However, small sample sizes have limited assessment of whether these mutations act as drivers (versus simply being observed by chance) in the cancers in which they occur more rarely. We reasoned that we could overcome this obstacle by analyzing large meta-cohorts of whole-exome sequencing and targeted sequencing samples, thus providing the statistical power to distinguish driver from passenger SNVs even in rarely-altered genes. Such analysis has previously been impeded by the exclusion of synonymous mutation calls from large, publicly available targeted sequencing datasets^57–61^, rendering the data incompatible with driver gene detection methods that rely on synonymous mutation counts to calibrate neutral mutation models. The mutation maps we generated allowed us to circumvent this difficulty.

We thus assembled a meta-cohort with nonsynonymous SNVs from 14,018 whole-exome and targeted-sequencing samples representing ten solid tumor types (median samples per cancer: 1,195; range: 515-3,110) (**table S13**) (**Methods**) and applied our mutation maps to explore the burden of known activating SNVs in oncogenes (obtained from the Cancer Genome Interpreter^62^) and pLoF SNVs in TSGs (**Methods**). For each cancer, we restricted our analysis to “long-tail” genes, which we defined as oncogenes and TSGs not associated with that cancer type in any of three recent, large pan-cancer surveys of driver genes^7,27,63^. All cancers carried a significant burden of activating SNVs in long-tail oncogenes (P<3.78×10^-9^ for all cohorts) (**Fig. 4a**, **fig. S17**, **table S14**), with 1.0-4.9% of samples (depending on the cancer) estimated to carry driver activating SNVs in long-tail oncogenes, an order of magnitude more than expected under the neutral mutation rate for each cancer (**Methods**). In long-tail tumor suppressor genes, the burden of pLoF SNVs was significant for all cancers except prostate cancer (P < 3.10×10^-4^ for each cancer but prostate; P=0.056 for prostate) (**Fig. 4b**, **fig. S17**, **table S15**), with 3.0-6.4% of samples inferred to carry a driver protein-truncating SNV in long-tail TSGs. These rates were consistent when we restricted the analysis to only whole-exome sequenced samples, though power to detect positive selection was decreased due to reduced sample size (**fig. S18**, **tables S16-S17**).

**Figure 4:**
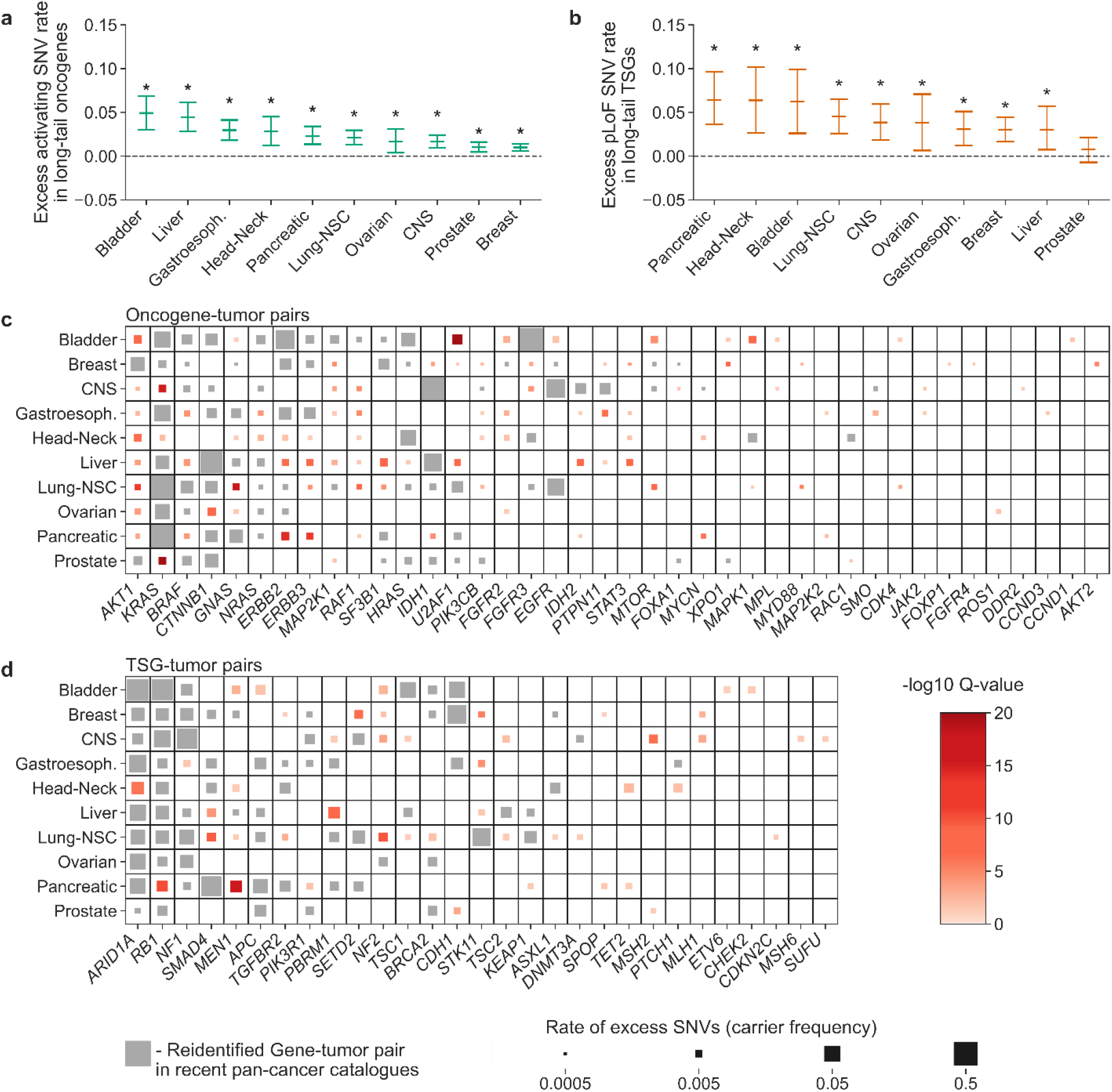
Enrichment of protein-altering SNVs in “long-tail” genes reveal a shared landscape of common and rare driver genes. **a,b,** Estimated rates of excess oncogenic SNVs in oncogenes (**a**) and predicted loss-of-function (pLoF) variants in TSGs (**b**) that were not previously associated with a given cancer (x-axis) in three large driver gene catalogues^7,8,63^. Stars indicate that the burden of oncogenic (pLoF) SNVs was significant in long-tail oncogenes (TSGs) in the cancer type. **c,d,** Oncogene-tumor pairs and TSG-tumor pairs with a significant burden of oncogenic or protein-truncating SNVs. Gene-tumor pairs previously reported by Dietlein et al.^7^, Bailey et al.^8^, or Martínez-Jiménez et al.^63^ are marked in grey. Pairs that are not present in those catalogues are marked in red with color intensity indicating significance of association. Marker size is proportional to the estimated rate of excess mutations after accounting for cancer-specific neutral mutation rates.

We also identified individual oncogenes and TSGs with a significant burden (FDR<0.1) of activating and pLoF SNVs, respectively (**Methods**). The increased sample sizes and targeted nature of the analysis enabled us to newly identify candidate driver genes with a median mutation frequency of 0.26% (interquartile range: 0.14-0.50%) compared to a median carrier frequency of 4.4% (interquartile range: 2.2%-9.1%) for driver genes in existing databases. Across the ten cancers, we identified 96 oncogene-tumor pairs not reported in recent pan-cancer surveys of driver genes (and reidentified 94 that were previously reported) (**Fig. 4c**, **table S18**). A control analysis of pLoF mutations in oncogenes found that only one of the 96 oncogene-tumor pairs also had a significant burden of truncating mutations (**fig. S19**). Similarly, we identified 46 TSG-tumor pairs not reported in the pan-cancer surveys (and reidentified 107 that were previously reported) (**Fig. 4d**, **table S19**). These results indicate that numerous driver mechanisms are shared across the common and rare driver landscape of solid cancers. For example, *KRAS* was significantly associated with all ten cancers (7 previously reported; 3 long-tail associations reported here) and *BRAF* was associated with nine of the ten cancer types (6 previously reported; 3 long-tail associations reported here). While some of these associations have been noted in clinical cases^56^, our results represent progress toward an unbiased, pan-cancer catalog of common and rare driver genes that we anticipate future studies will further elucidate.

## Discussion

Our study demonstrates that the ability to analyze specific, carefully constructed combinations of mutations anywhere in the genome for evidence of selection can empower the identification of candidate cancer drivers. To enable such an analysis, we introduced a method that estimates the mutation rate genome-wide through a combination of deep-learning and probabilistic modeling. The strong performance of the method in modeling mutation rates and identifying candidate drivers highlights how deep-learning algorithms can empower interpretable biological discovery by accurately estimating key modeling parameters (**Note S2**). Leveraging the flexibility of this approach, we identified candidate driver elements with a significant enrichment of mutations in understudied regions of the genome. While the coding and noncoding driver candidates we report – in cryptic splice sites, 5’ UTRs, and “long-tail” genes – occurred at low frequencies individually, our estimates suggest that they collectively contribute to the disease pathology of up to 10% of tumors. Moreover, this estimate may be conservative, as several of our analyses utilized datasets of mutations that are unlikely to be comprehensive (e.g., catalogs of predicted cryptic splice mutations and known activating missense mutations).

These aggregate numbers have potential clinical ramifications. First, they emphasize the utility of profiling the mutational landscape of clinical tumor samples either with whole genome sequencing or broad targeted sequencing capable of capturing driver mutations in diverse regions of the genome. While some cancer centers have started to sequence each tumor using panels of hundreds of known driver genes^56^, our results hint that further expanding these panels to include some intronic and untranslated sequence may yield additional mutations of possible clinical significance. They also suggest potential therapeutic strategies, for example oligonucleotides to mitigate cryptic splice mutations^64–66^. Of particular interest may be *TP53* 5’ UTR splice mutation carriers; since these mutations were mutually exclusive to coding mutations in *TP53,* their *TP53* transcripts likely encode a fully functional p53 protein that could potentially be translated upon suppression of incorrect splicing within the 5’ UTR.

Our results provide a few glimpses into the landscape of noncoding driver mutations that we anticipate will emerge as cancer sequence sample sizes continue to grow. A power analysis we performed suggested that current sample sizes are not adequate to uncover infrequent drivers under moderate or weak positive selection (**fig. S20**; **Methods**), which have been proposed to underlie a polygenic “mini driver” model of cancer evolution^67^. We anticipate that our maps will be particularly useful in evaluating this and other hypotheses due to their ability to consider mutations spread over large swaths of the genome. For instance, a preliminary analysis we performed of enhancer networks identified several genes with a burden of enhancer mutations (**table S20, Note S5**), including *FOXA1,* in which promoter mutations are thought to drive breast cancer by increasing gene expression^68^.

However, care is warranted in interpreting the potential driver role of any locus deemed to carry a burden of mutations, particular in the absence of functional data. While an excess of mutations compared to the neutral mutation rate provides evidence in support of positive selection, it does not prove selection because no neutral mutation model is perfect; thus, mutational excess can reflect model inaccuracies instead of bona fide positive selection. While our approach of explicitly modeling mutational patterns at the kilobase and nucleotide scales captured, on average, fine-scale mutation rates across the genome, accounting for local, likely-neutral hypermutation processes observed at particular focal loci^5,17^ remains a challenge and an area for future innovation.

## Methods

Detailed methods for this manuscript have been provided as supplementary material.

## Data availability

Data generated as part of this study are available as supplementary tables or from http://cb.csail.mit.edu/cb/DIG/. Browsable mutation maps for 37 cancer types are provided at https://resgen.io/maxsh/Cancer_Mutation_Maps/views. PCAWG data are available from https://dcc.icgc.org/releases/PCAWG/. Hartwig Medical Foundation data are available from https://database.hartwigmedicalfoundation.nl/. Whole exome sequencing data compiled by Dietlein et al. are available from http://www.cancer-genes.org/. Targeted sequencing data are available from https://www.cbioportal.org/.

## Code availability

The method described in this manuscript (Dig) is available as a package hosted on the conda repository (https://anaconda.org/mutation_density/digdriver). Installation instructions and documentation are available at https://github.com/maxwellsh/DIGDriver/wiki. The Dig source code is available on GitHub (https://github.com/maxwellsh/DIGDriver). All other code used in this study are available from the authors upon request.

## Acknowledgements

We thank A. Barton, M. Hujoel, R. Mukamel, L. Sholl, and P. Park for their valuable comments and suggestions. We thank I. Martincorena and F. Abascal for graciously providing source code and instructions for running NBR. The results presented in this study are based in part on data generated by the ICGC and TCGA PCAWG Network, the TCGA Research Network, the Hartwig Medical Foundation, and the Memorial Sloan Kettering Cancer Center. M.A.S. was funded in part by NIH fellowship F31 MH124393. F.D. was supported by the Claudia Adams Barr Program for Innovative Cancer Research and an ASPIRE Award of The Mark Foundation for Cancer Research. P.-R.L. was supported by NIH grant DP2 ES030554, a Burroughs Wellcome Fund Career Award at the Scientific Interfaces, the Next Generation Fund at the Broad Institute of MIT and Harvard, and a Sloan Research Fellowship. B.B. was supported in part by NIH grant U01CA250554 from the National Cancer Institute.

## Contributions

M.A.S., A.Y., and B.B. designed the Dig method. M.A.S., A.Y., and O.P. implemented software code. All authors conceived and designed analyses. M.A.S., A.Y., and O.P. performed the analyses. All authors wrote the manuscript and prepared figures.

## Ethics declaration

M.A.S, A.Y. and B.B. are co-inventors on a provisional patent related to the Dig method.

