## Supplementary material for "Learning the mutational landscape of the cancer genome"

### Methods

#### Datasets of somatic SNVs and indels

*PCAWG dataset:*

We obtained somatic SNVs and indels from whole-genome sequencing of 2,955 unique tumor samples across 38 types of cancer from the ICGC data portal (https://dcc.icgc.org/) and dbGaP (project code: phs000178). We restricted to a set of 2,583 samples that had previously passed quality control checks[1] and constructed 14 meta-cohorts that combined similar tumor types as described in Rheinbay et al. Consistent with Rheinbay et al., our pan-cancer meta-cohort excluded melanoma and hematopoietic malignancies. We removed samples with reported high microsatellite instability from each cohort and meta-cohort with the exception of the pan-cancer cohort. We then annotated autosomal SNVs and indels with their predicted coding impact (synonymous, missense, nonsense, essential splice) using a custom annotation method (Code Availability) that we adapted from the dNdScv software[2]. (We excluded sex chromosomes from this study due to the differing X and Y mutation rates between males and females). For the creation of reference neutral mutation models and driver element analysis, we considered 37 cancer cohorts and meta-cohorts composed of at least 20 samples and $10^5$ SNVs (**table S2**). The annotated mutations from the ICGC portion of the PCAWG dataset are publicly available on our website (**Data Availability**), and we provide instructions for generating the TCGA portion of the dataset for researchers with approved TCGA access.

*Dietlein et al. dataset:*

We obtained somatic SNVs and indels from whole-exome sequencing of 8,617 tumor samples across 17 cancer types that had previously been curated in Dietlein et al.[3] (http://www.cancer-genes.org/) for which the same cancer type was also assayed by PCAWG (**table S21**). We additionally constructed a pan-cancer dataset by merging somatic mutations from all samples excluding melanoma and hematopoietic malignancies as above. Mutations were annotated for their function in coding sequence as described above. The annotated mutations are publicly available on our website (Data Availability).

*Target sequencing datasets:*

We obtained somatic SNVs from targeted sequencing of 10 types of solid cancers performed using the IMPACT protocol at the Memorial Sloan Kettering Cancer Institute from cbioportal [4] (https://www.cbioportal.org/) (**table S13**). We removed duplicate samples and hypermutated samples with >100 coding mutations in 221 genes common to all whole-exome and targeted sequenced samples. SNVs were then annotated for their function in coding sequence as described above and merged with SNVs from the whole-exome datasets (after removing hypermutated samples) to form mega-cohorts with aggregate sample size of 14,018 tumors. The annotated mutations are publicly available on our website (**Data Availability**).

#### Learning kilobase-scale somatic mutation rates genome-wide

Our approach (Dig) leverages two key insights from cancer genomics: 1) while the passenger mutation rate varies dramatically across the cancer genome[5,6], it is approximately constant at the kilobase scale [1]; and 2) the pattern of passenger mutations within a region is highly dependent on the nucleotide context of each base[3,7–9]. For each cancer, our mutation "maps" thus consists of two components: a "region model" providing the mean and variance of the passenger mutation count in kilobase-scale regions tiled across the genome and a "sequence context model" that provides the genome-wide likelihood of a mutation in any given nucleotide context. These components are then integrated via a probabilistic model to estimate the neutral mutation rate in any genomic element. Intuitively, the region model provides information about how many neutral mutations should be present in a region while the sequence context model determines how many of those mutations should be distributed to any sub-element of the region. Crucially, the components of the model for a given cancer need to be estimated only once; they can then be publicly distributed and utilized by anyone for driver discovery in a dataset of interest. Each element of the model is described in detail below.

*Input data for model training*

Training the mutation map model requires three datasets as input: 1) A dataset of somatic SNVs from a whole-genome sequenced cohort of tumors; 2) high-resolution epigenetic assays; and 3) the human reference genome corresponding to the build used for calling the somatic SNVs and aligning the epigenetic assays (the build must be the same for the SNV and epigenetic datasets). In this study, we used somatic SNVs from the PCAWG dataset (described above). For epigenetic tracks, we obtained $-\log_{10}$ P-values genome-wide for peak calls for all "reference epigenomes" of healthy tissues uniformly processed by the Roadmap Epigenomics consortium[10] (127 epigenomes totaling 723 chromatin assays) and replication timing assays of 10 cell lines from ENCODE[11] (additional details and download links in **table S1**); we additionally constructed two custom tracks quantifying the average nucleotide and GC content in every non-overlapping 100bp window in the genome.  For the human reference, we used the hg19 / GRCh37 build, which was used by both PCAWG and Roadmap in the generation of their datasets. We have made the epigenetics datasets in formats compatible with our software and the reference genome used in this study publicly available (**Data Availability**).

*Estimating mutation rates in kilobase-scale regions*

We estimated mutation rates for each cancer cohort in non-overlapping windows of 10kb tiled across the genome using a probabilistic deep learning model consisting of a composition of a convolution neural network (CNN) and Gaussian process (GP) (**Fig. 1a**, **fig. S1b**). This composite model had the desirable property of both predicting the expected mutation rate and simultaneously quantifying the prediction uncertainty, a property that has been shown to improve out-of-sample prediction[12] and is essential to our probabilistic refinement of mutation rates (described below).

The CNN took as input matrices of size $735 \times 100$ where each row is an epigenetic feature track (describe above) and each column is the average track value in non-overlapping 100bp windows. Optionally, the CNN also took as input the observed SNV counts in 100kb regions flanking each 10kb region in the training SNV dataset. The CNN architecture is as follows: it contains 4 convolutional blocks with 2 batch normalized convolutional layers and ReLU activation. The first block reduces the $735 \times 100$ input tensor to $256 \times 50$ with 256 channels and a double stride. The following blocks are ResNet-style residual blocks which maintain their input dimension to facilitate residual connections with 256, 512, and 1024 channels respectively. Between each of the 3 residual blocks there is a double stride (ReLU activated and batch normalized) convolutional layer, which reduces the tensor length by half and doubles its height with additional channels. The output of the last residual block is flattened (and optionally concatenated with the two-flanking region counts) and passed through 3 fully connected (FC) layers. The first two FC layers are ReLU activated and reduce the dimensionality of the vector to 128 and 16 dimensions respectively. The last FC layer performs the final regression that predicts the SNV count in the 10kb region via a linear function. The CNN architecture was implemented in PyTorch[13]. The Gaussian process is a sparse, inducing-point GP[14] with a radial basis function kernel that takes as input the final 16-dimensional feature vector of the trained CNN and non-linearly predicts both the mean and variance of the neutral mutations in the associated 10kb region. The GP architecture was implemented in GPyTorch[15].

Genome-wide prediction of the mutation rate in 10kb windows was accomplished via a 5-fold cross-validation scheme that rigorously ensured that the training input data was separate from the windows over which the mutation rate was being predicted. We first removed regions likely to contain spurious mutation counts, defined as windows where less than 50% of the

36mers uniquely mapped back to that region or regions in the top 99.99th percentile of mutation counts. The remaining 10kb windows were then divided into five non-overlapping sets, each containing 20% of the windows. Sequentially, each set of windows was held-out, a new deep-learning model was trained over the remaining four window sets (80% of the genome), and prediction was preformed over the held-out set (20% of the genome); prediction was also performed over the removed regions. Thus, mutation rates for each 10kb window in the genome was predicting using a model that had not been trained on that window.

The CNN and GP were trained sequentially. First, the CNN was trained for 20 epochs with a batch size of 128 samples, using the Adam optimizer to minimize mean squared error loss to the observed mutation counts in each training window. For training, the input data was additionally divided via an 80-20 split into training data and validation data (thus for each fold, 64% of the genome was used for training, 16% for validation, and 20% for held-out prediction). To avoid overfitting the data, the epoch from which the trained CNN was selected was determined by highest validation R-squared accuracy to observed counts in the validation-set across all CNN epochs. Once the CNN was trained, the final 16-dimension feature vector for each training window was passed as input to the Gaussian process which was trained to predict the observed mutation counts in each training window by minimizing a multivariate normal loss function with the Adam optimizer. The GP was optimized with 400 inducing points for 50 iterations. Due to the inherent variability in gradient-based optimization, we ran the GP 10 independent times and calculated the ensemble average of the mean and variance predictions from each of the individual runs. These ensemble predictions were then used as the mean and variance estimates for each 10kb region.

Some random initializations of the GP would fail to converge (defined as a decrease in R-squared accuracy of more than 0.03 compared to the final accuracy of the trained CNN). When this occurred, the GP was restarted up to 3 times to achieve a successful convergence. If after 3 attempts, the GP had not successfully converged, the number of inducing points was reduced by 100 and the GP given another 3 attempts to converge. This process continued until successful convergence or a reduction to zero inducing points. If a GP failed to converge in all 12 attempts, the CNN was reinitialized to generate a new set of feature vectors.

*Training the sequence-context model*

The nucleotide-level mutation rate is heavily influenced by the trinucleotide context of a position[16,17]. For each cancer, we thus estimated the likelihood that each of 192 possible trinucleotide mutations could occur (four possible upstream nucleotides by four possible middle nucleotides by three possible mutations by four possible downstream nucleotides) using a maximum-likelihood estimation approach. Specifically, let $V_{aX \to Yb}$ be the number of times that the nucleotide context $a, X, b$ is observed with $X$ mutated to the nucleotide $Y$ in the training cancer cohort and let $N_{aXb}$ be the number of times the trinucleotide $a, X, b$ occurs in the genome, where $a, X, b \in \{A, C, T, G\}$ and $Y \in \{A, C, T, G\} \setminus X$. Then the genome-wide likelihood of the mutation $aX \to Yb$ was estimated as $\Pr(aX \to Yb) = p_{aX \to Yb} = \frac{V_{aX \to Yb}}{n \cdot N_{aXb}}$ where $n$ is the total number of samples in the training cohort. In practice, we do not need to divide each estimate by $n$ because the probabilities are normalized to sum to one during the refinement step (see below), so the constant is extraneous. While we chose to use a trinucleotide model in this work, any model that predicts the likelihood of a mutation solely from sequence context could be used, for example a pentanucleotide model or more complex composite likelihood model[3].

#### Refining kilobase-scale mutation rate estimates to test for evidence of selection at the mutation scale

We derived a probabilistic method to estimate a distribution over the number of SNVs and indels observed at a set of positions in a dataset of interest given the kilobase-scale estimated mutation rate $\mu_R$ and estimation uncertainty $\sigma_R^2$ along with the sequence context likelihood estimates. We refer to this method as Dig.

*Passenger model for SNVs*

Let $M_{i,aX\to Yb}$ be the (discrete) number of mutations of the form $aX \to Yb$ at position $i$; $X_R$ be the number of mutations in a kilobase-scale region $R$; and $\lambda_R$ be the *rate* at which mutations occur in the region $R$. We assume the following generative model for how mutations accumulate within the genome for a cancer of interest (**fig. S22**):

$$\Pr\left(M_{i,aX\to Yb} = k, X_R, \lambda_R\right) = \Pr\left(M_{i,aX\to Yb} = k \mid X_R\right) \cdot \Pr(X_R|\lambda_R) \cdot \Pr(\lambda_R)$$

where

$$\lambda_R \sim \mathrm{Gamma}(\alpha_R, \theta_R)$$

$$X_R|\lambda_R \sim \mathrm{Poisson}(\lambda_R)$$

$$M_{i,aX\to Yb}|X_R \sim \mathrm{Binomial}\left(X_R, p_{R,aX\to Yb}\right).$$

The parameters $\alpha_R$ and $\theta_R$ are the shape and scale parameters of a gamma distribution, respectively. The parameter $p_{R,aX\to Yb}$ is the normalization of $p_{aX\to Yb}$ such that the probability of all possible mutations in $R$ sums to one.

While the value of $M_{i,aX\to Yb}$ is observed in a cancer cohort of interest, $X_R$ and $\lambda_R$ are unknown parameters, and thus must be integrated out for the model to be of practical use. We showed previously that the generative model is an extension of the classic Poisson-Gamma distribution *and* that the marginal distribution $\Pr\left(M_{i,aX\to Yb} = k\right)$ has a simple closed form[18]:

$$M_{i,aX\to Yb} \sim \mathrm{NegativeBinomial}\left(\alpha_R, \frac{1}{1 + \theta_R \cdot p_{R,aX\to Yb}}\right).$$

Moreover, assuming that SNVs arise approximately independently, this extends to any set of SNVs $I \subseteq R$ as

$$\sum_I M_{i,aX\to Yb} \sim \mathrm{NegativeBinomial}\left(\alpha_R, \frac{1}{1 + \theta_R \cdot \sum_I p_{R,aX\to Yb}}\right).$$

We employ a variational approach to estimate $\alpha_R$ and $\theta_R$. In particular, values for these two parameters uniquely determine the mean and variance of a gamma distribution. Thus, mean and variance estimates for a region $R$ uniquely determine the values for $\alpha_R$ and $\theta_R$. Given $\mu_R$ and $\sigma_R^2$ estimated by our deep-learning method, the variational estimates for $\alpha_R$ and $\theta_R$ are:

$$\alpha_R = \frac{\mu_R^2}{\sigma_R^2}$$

$$\theta_R = \frac{\sigma_R^2}{\mu_R}.$$

Finally, since $\mu_R$ and $\sigma_R^2$ are estimated for a particular cancer cohort used for training, they must be updated to account for any differences between the training cohort and the target cohort of interest. Assuming that the mutational processes (and technical analysis of the training cohort and target cohort) are similar, the model for the training cohort can be readily adjusted to the target cohort by the introduction of a single scaling factor $C_{\text{SNV}}$ that, intuitively, accounts for the difference in sample size between the training and target cohorts. Under this formulation, the distribution over a set of possible SNVs in a region is:

$$\sum_I M_{i,aX\to Yb} \sim \text{NegativeBinomial}\left(\alpha_R, \frac{1}{1 + C_{\text{SNV}} \cdot \theta_R \cdot \sum_I p_{R,aX\to Yb}}\right).$$

By default, we estimate the scaling factor as the ratio of the number of observed synonymous SNVs in the target dataset to the number of expected synonymous SNVs in the training cohort across all genes excluding *TP53* (in which some synonymous mutations are under positive selection [2]). However, $C_{\text{SNV}}$ can be equivalently estimated by other approaches when synonymous mutations are not available or are observed in only some genes (see below and **fig. S18**).

The assumption of biological and technical similarity between the training and target cohort introduces an important limitation to our approach. The training and target cohorts must be carefully matched, similar to how genetic ancestry must be matched between an imputation panel and a target SNP dataset in population genetics in order for imputation of genetic variation to be accurate. The standardization of sequencing and variant calling pipelines in recent years likely minimizes the extent of technical differences between datasets. However, matching the type of cancer between the training and target cohort is crucial.

*Passenger model for indels and multi-nucleotide variants*

The indel model is identical to that of the SNV model with two exceptions. First, we assume a uniform distribution of indels independent of sequence context, as has been assumed in previous works[2]. Thus in the negative binomial distribution above, $\sum_I p_{R,aX\rightarrow Yb}$ is replaced by the uniform mutation probability $|I|/|R|$ where $|\cdot|$ denotes the total number of genomics position in $I$ and $R$. The uniform assumption of indels could readily be replaced with a probability distribution based on indel type, size and homology [9], but we do not pursue that extension here. Second, the scaling factor for indels, $C_{\text{indel}}$, is estimated as the ratio of the number of indels observed in the target dataset to the number of expected indels in the training dataset across the coding sequence of all genes not in the Cancer Gene Census. We treat multi-nucleotide variants (MNVs) as indels.

We tested estimating $\mu_R$ and $\sigma_R^2$ independently for SNVs and indels using separate deep learning models for the two types of mutations. We found that direct estimation of these parameters for indels resulted in a less accurate indel model than using the SNV estimates as a proxy for indel estimates. We suspect this is due to the fact that indels occur an order of magnitude less frequently than SNVs and thus there are too few observed indels in the training cohort for the deep-learning model to build an accurate prediction function. As sample sizes become larger, we expect that directly training a deep-learning model to predict indels will yield more accurate predictions.

*Extension to mutations spanning multiple kilobase-scale regions*

We take two approaches to extend the above passenger models to account for sets of mutations that span multiple kilobase-scale regions.

- Approach 1: approximate the distribution across the regions by extending the variational estimation of $\alpha_R$ and $\theta_R$. Specifically, let $R' = \{R_1, \dots, R_n\}$ be the set of regions in which a set of mutations occur. Then we estimate $\mu_{R'} = \sum_{i=1}^{n} \mu_R$ and $\sigma_{R'}^2 = \sum_{i=1}^{n} \sigma_R^2$, and $\alpha_{R'}$ and $\theta_{R'}$ are then estimated as above from $\mu_{R'}$ and $\sigma_{R'}^2$.
- Approach 2: exactly estimate the distribution across the mutation set by convolving the distributions arising from the subset of mutations in each $R_i \in R'$.

Approach 1 is computationally efficient and accurate so long as the mutation rate estimates across $\{R_1, \dots, R_n\}$ are sufficiently similar. Thus approach 1 is preferred when $R'$ is composed of a small number of contiguous (or nearly contiguous) regions and is the default implemented algorithm. When $R'$ is composed of regions with highly variable mutation rates, approach 1 is likely to either over- or under-estimate that passenger mutation rate, leading to improperly calibrated p-values. In this case, approach 2 will provide accurate estimates but requires more computation due to need to perform convolution.

*Testing mutational burden across a set of candidate mutations using an existing mutation map*

The steps to estimate selection using Dig are as follows:

User steps:

1. Download a mutation map for the cancer matching the cancer type of the dataset of interest.
2. Provide the mutation dataset of interest and define the set $I$ of possible mutations. $I$ can be defined as any set of genomic intervals (contiguous or noncontiguous) or any set of possible SNVs anywhere in the genome.

Software steps:

3. The mutation likelihoods $p_{R,aX\rightarrow Y,b}$ and $p_{\mathrm{indel}}$ are calculated as described above for each mutation set. The nucleotide sequence of $R$ is extracted from the reference genome.
4. The SNV and indel scaling factors are estimated for the cohort of interest.
5. The p-value of the number of SNVs and indels observed in the cohort of interest for each mutation set are calculated using the negative binomial distributions defined above as the null models.
6. The p-values for the SNVs and indels are combined via Fisher's method.

For this study, we used the mutation maps trained using both epigenetic tracks and flanking mutation counts to test for burdens of mutations. These are also the maps we have made publicly available.

#### Comparison of passenger mutation model to existing methods

We assessed the variance explained of observed SNV counts by the estimated mutation counts produced by our approach (Dig) across diverse genomic elements. We compared this to the variance explained by existing methods that construct passenger mutation models. Variance explained was quantified as the square of the Pearson correlation coefficient between the observed and predicted mutation counts.

*Tiled regions*

We compared the variance explained (square of the Pearson correlation coefficient) in SNV counts within 10kb windows tiled across the genome between Dig and NBR [1]. NBR is, to our knowledge, the only method that has been previously used to build passenger mutation models in kilobase-scale regions tiled across the genome. However, code for running the NBR method is not currently publicly available. For each cancer, the NBR model was trained on the same regions used to train our deep-learning model (excluded regions with 36mer mappability <50% and regions in the top 99.99$^{th}$ percentile of mutation count). The regions excluded from training were also excluded when calculating the variance explained statistic.

In discrete stochastic systems, random stochasticity of events when event rate is low results in deflation of the variance explained statistic. Intuitively, for a discrete process with event rate <1, the expected value will be a real value between zero and one but the observed count will be a whole number (likely either zero or one); thus even a method that perfectly predicts the expected value will still have a low variance explained statistic because the observed values will almost always deviate substantially from the expected value. To guard against this deflation, we required that, on average, the expected number of mutations per 10kb region be approximately one (thus ensuring that the standard deviation was of the same order of magnitude or less than the expected value). Thus, in *Results*, variance explained statistics in 10kb regions were reported only for cancer cohorts with >1 million mutations. Variance explained for all cancer types are reported in **table S3**.

We also assessed the variance explained of SNV counts in 1Mb regions by our method (restricted to 1Mb regions with >50% 36mer mappability). To estimate the expected mutation count in each 1Mb region, we summed together the estimates of each non-overlapping 10kb window within the 1Mb region.

*Coding sequence*

We compared the variance explained in *nonsynonymous* SNV counts between Dig and two widely used methods that generate nonsynomous SNV passenger mutation models: MutSigCV[5] and dNdScv[2]. Both MutSigCV and dNdScv utilize the synonymous mutations observed in each gene to estimated gene-specific passenger mutation rates. Variance explained was evaluated over the coding sequence of 3,740 genes that were 1) common to all three methods; 2) between 1kb and 1.5kb in length; and 3) not in the CGC. The length restriction was imposed to prevent coding sequence length from artificially inflating variance explained since the number of mutations in a gene strongly correlates with its length.

*Noncoding regulatory elements*

We compared the variance explained in SNV counts between Dig and two other methods that estimate passenger mutation rates in noncoding regulatory elements: DriverPower[19] and Larva[20]. DriverPower is optimized to estimate mutation rate within a set of regulatory elements predefined by the authors of the software; this set of elements is not easily changed. We thus evaluated variance explained in a set of 7,412 noncoding regulatory elements (enhancers, lncRNAs, and scRNAs) between 0.5kb and 1kb in length that could be modeled by DriverPower. The length restriction was again implemented to prevent inflation of variance explained due to variance in element length. While Larva can predict mutation rate within genomic intervals, it cannot natively provide a prediction for elements that are composed of multiple, non-contiguous intervals. To circumvent this, we divided each element evaluated by DriverPower into its constituent intervals, produced a prediction for each interval separately with Larva, and summed the predictions across regions composing a single element.

#### Comparison of driver detection accuracy to existing models

*Coding sequence models*

We compared the sensitivity, specificity, and F1-score (harmonic mean of sensitivity and specificity) for driver gene detection from coding sequence mutations between Dig, MutSigCV, and dNdScv across the 37 PCAWG cancer cohorts and 17 whole-exome sequenced cohorts. We chose to compare to these two methods because they are widely used driver gene detection methods that rely on neutral mutation models to test for selection. An FDR significance threshold of 0.1 was applied for all methods and cohorts. A true-positive driver gene was defined as any gene in the Cancer Gene Census (CGC)[21] that was detected as FDR significant by any of the methods in a given cohort. A false-positive was defined as any gene identified as FDR significant that was not in the CGC. Each method was applied to the same set of 16,794 genes. Both SNVs and indels were used to identify potential driver genes. Our approach and dNdScv were compared over two sample filtration regimes: 1) no samples filtered and 2) samples filtered by dNdScv's default filtration settings (samples with >3000 coding mutations excluded and total mutations per gene per sample capped at 3). MutSigCV does not offer user-definable filtration settings. All methods were otherwise run with default settings.

*Noncoding sequence models*

We compared the sensitivity, specificity, and F1-score for driver noncoding element identification from noncoding SNVs between Dig, DriverPower, Larva, and ActiveDriverWGS[22] across the 37 PCAWG cancer cohorts. We chose to compare to these three methods because they are recently introduced methods for noncoding driver element identification that rely on neutral mutation models to test for selection. An FDR significance threshold of 0.1 was applied for all methods and cohorts. A true-positive driver element was defined as any element previously identified by PCAWG as carrying a burden of mutations[1] that was detected as FDR significant by any of the methods in a given cohort. A false-positive was considered any FDR significant element that was not previously identified by PCAWG as having a burden of mutations. This comparison was conservative (biased against our approach) for two reasons: 1) The other three methods were previously applied to the PCAWG dataset to generate the set of putative driver elements that we then used as a gold standard for the same samples; and 2) we restricted the analysis to SNVs because not all methods we compared to could accept indels. Indeed, our approach is the only approach that models SNVs and indels independently; the other approaches either do not model indels or model indels and SNVs as a single category.

#### Associating epigenetic structure to mutation density with feature maps

To investigate the underlying features the deep learning model considered when predicting mutation rates, we added another layer of computation between the input epigenetic matrix and the CNN to serve as feature maps. Feature maps are a tool used in computer vision tasks to detect which regions of an image the model uses to perform prediction[23]. We used this technique to evaluate which epigenetic patterns the CNN exploited to predict mutation rates. To reduce the potential for noise, we applied this technique to input matrices encoding 50kb regions.

*Feature map generation*

An additional two-layered network was added between the input matrix and CNN to force the model to attend to the subset of most salient input sub-regions and compute the feature maps. In the attention augmented CNN, the input matrix was first passed through two convolutional layers preserving the input dimensionality (stride length 1, kernel sizes 5 and then 3) with ReLU activations. Subsequently, the output of the two layers was passed through a row-wise Softmax function that had the effect of making most entries in the matrix close to zero with sparse values close to one. The resulted “feature map” matrix was then element wise multiplied with the original input and passed on to the downstream CNN. This had the effect of setting most entries in the original epigenetic matrix to near zero, thus forcing the CNN to rely only on the small subspace of the input that was not zeroed out. The optimization process compels the feature maps to attend to the features of the input matrix most relevant for the prediction process.

*Extraction of epigenetic content of feature maps via dimensionality reduction and clustering*

While the feature maps have the theoretical ability to attend to any regions of the input matrix, in practice we found they almost always attended to a large set of epigenomic features (rows) in a small set of contiguous columns (genomic positions), zeroing out most values outside of these columns. We extracted and summarized the epigenetic content of each these attention columns through the following approach: 1) in each 50kb window, we extracted the largest contiguous set of columns such that each column contained at least 10 cells with a non-zero entry. This contiguous set of columns was defined as an “attention super-column”. 2) Each attention super-column was reduced to an 8-dimensional vector by averaging together tracks of the same epigenetic type per column (DNase, H3K27ac, H3K27me3, H3K36me3, H3K4me1, H3K4me3, H3K9ac, and H3K9me3) and taking the maximum value across each row. 3) The vectors were normalized and projected into a two-dimensional subspace for clustering and visualization via a Uniform Manifold Approximation and Projection (UMAP) transformation (Python UMAP-learn package; #neighbors: 30, minimum distance: 0, #components: 2). 4) Spectral clustering (Python Sklearn package) was applied to the two-dimensional subspace to identify attention super-columns with similar epigenetic content. The clustering consistently identified five distinct clusters across cancer cohorts.

*Connecting feature map epigenetic clusters to functional annotations*

To determine whether the attention super-columns in a cluster represented a functional epigenetic structure, we extracted the average Epilogos[24] signature vector per attention super-column and examined whether the Epilogos signatures were consistent within a cluster. Epilogos is a summary of the functional epigenetic states across 111 tissue types as inferred by the ChromHMM method.

*Connecting feature map epigenetic clusters to mutation rates*

We extracted the mutation count for the 50kb window in which each attention column occurred. We computed the mean and standard deviation of mutation counts across the attention column clusters.

#### Constructing a genome-browser of genome-wide mutation rate estimates

We used Dig to estimate mutations rates in every non-overlapping regions of size 100bp, 250bp, 500bp, 1kb, 2.5kb, 25kb, 50kb, 100kb, 250kb, 500kb and 1Mb tiled across the genome (excluding assembly gaps in the GRCh37 reference genome) for 37 PCAWG cancer types. These predictions were used to construct data structures that can be interactively visualized by HiGlass[25].

#### Quantifying selection on cryptic splice SNVs

*Curation of predicted splice SNVs*

We obtained predictions for the splicing impact of every possible SNV in the body of 17,816 autosomal genes from SpliceAI[26]. We restricted to SNVs with predicted splicing impact (i.e., SpliceAI Δ score) >0.2 and separated canonical from cryptic splice SNVs. A canonical splice SNV was defined as an SNV altering positions 1 or 2 base-pairs 5' or 3' to an exon boundary of any transcript in the Gencode V24 list of basic transcripts. Cryptic splice SNVs were defined as all other SNVs with Δ score >0.2, excluding sites that were 5 base-pairs 3' to an exon boundary that had been included in the definition of "essential splice sites" considered by Marticorena et al. [2]. (We excluded these sites to ensure that any enrichment we observed in mutation rates at cryptic splice sites were independent of enrichment previously noted by Martincorena et al.). In sub-dividing cryptic splice SNVs into coding and noncoding, we defined coding SNVs to be *synonymous* SNVs falling within the canonical transcript of each gene and noncoding SNVs to be those falling outside of the coding sequence of *all* Gencode V24 basic transcripts for each gene.

*Quantifying enrichment of coding mutations and splice SNVs in the PCAWG dataset*

For each gene, we applied Dig with default settings to map the mutation rate across the following sets of mutations in the PCAWG pan-cancer dataset: synonymous SNVs, missense SNVs, nonsense (stop-gained) SNVs, coding indels, canonical splice SNVs, cryptic splice SNVs (stratified by predicted splicing impact into low ($0.2 \leq \Delta < 0.5$), medium ($0.5 \leq \Delta < 0.8$) and high ($0.8 \leq \Delta < 1.0$) impact and by coding versus noncoding designation). Across a set of genes (e.g., TSGs, oncogenes, genes not in the CGC), we defined mutation enrichment for a given mutation type to be the ratio of the observed mutations to expected mutations summed across the gene set. This statistic is conceptually similar to the selection coefficient reported for coding mutations by dNdScv.

P-values for a gene set and mutation type were exactly calculated by convolving the mutation-type specific negative binomial distributions for each gene in the gene set and summing the upper-tail probability that at least the number of observed mutations occurred by chance. We used the following Monte Carlo simulation approach to estimate the 95% confidence intervals of enrichment within a gene set and mutation type.

1. For each gene, estimate the enrichment coefficient as the number of observed mutations divided by the number of expected mutations. A small pseudo-count of $1 \times 10^{-16}$ was added to the numerator and denominator to prevent the enrichment from being identically zero when no mutations were observed in a gene. (This would lead to a degenerate Poisson distribution in step 3).
2. For each gene, randomly draw a Poisson rate parameter from the gamma distribution defined by the mean and standard deviation estimates of the kilobase-scale mutation rate map.

3. For each gene, randomly draw a number of “observed” mutations from a Poisson distribution with rate parameter equal to the simulated rate parameter multiplied by the enrichment coefficient and the likelihood of the mutation type occurring within the gene. Conceptually, this mutation count is simulated under the hypothesis of *positive selection* on the mutations within the gene.
4. Estimate a simulated enrichment by summing the number of simulated mutations across all genes in the set and dividing by the expected number of mutations under the null model of no enrichment.
5. Repeat steps 1-4 one thousand times and define the boundaries of the 95% confidence interval as the lower 2.5$^{th}$ percentile and upper 97.5$^{th}$ percentile of the simulated enrichments.

We quantified enrichment in all samples in the pan-cancer cohort (**Fig. 2b**) and excluding hypermutated samples with >3000 coding mutations (**fig. S12**).

*Additional quantification of mutation enrichment in TSGs and oncogenes*

To gain additional confidence in the accuracy of our mutation enrichment estimates, we directly compared the mutation rate in genes not in the CGC to TSGs and oncogenes in the CGC using a Fisher’s exact test. This approach recapitulated the enrichment patterns we observed using Dig. However, the Fisher’s exact test does not account for global mutation rate differences between genes not in the CGC and genes in the CGC; thus, the precise estimates in **fig. S13** are unlikely to be accurate.

*Identification of individual TSGs enriched for noncanonical cryptic splice SNVs*

In each of the 37 PCAWG cohorts, we identified TSGs in the CGC with a significant burden of noncanonical cryptic splice SNVs under the null model estimated by our method. The significance threshold was defined per cancer as FDR q-value<0.1 corrected for the number of tested TSGs (n=283). We excluded one significant gene, *PRDM1*, from further analysis because the observed excess mutations were attributable to a single sample.

*Enrichment of predicted splicing impact in noncoding cryptic splice SNVs observed in significantly burdened TSGs*

We used a bootstrap method to calculate a p-value for the null hypothesis that noncanonical cryptic splice SNVs observed in the genes with a significant burden of cryptic splice SNVs had a predicted impact on splicing similar to the predicted impact of cryptic splice SNVs observed in genes not in the CGC. We calculated the median of the Δ scores randomly resampled from the observed cryptic splice SNVs in the TSGs and observed cryptic splice SNVs in genes not in the CGC ten thousand times (the number of SNVs sampled from the non-CGC set was equal to the number observed in the TSG set). We estimated the p-value as the number of times the resampled median of the non-CGC cryptic splice SNVs exceeded the resampled median of the cryptic splice SNVs observed in the TSGs.

*Quantification of the pan-cancer contribution of cryptic splice SNVs to TSG driver SNVs*

We calculated the excess of SNVs in TSGs in the CGC stratified by function (missense, nonsense, canonical splice, and noncoding canonical splice) as the difference between the number of mutations observed and the number expected. The relative contribution for each category was defined as the excess for that category normalized by the sum of the excess across all categories. The 95% confidence interval for the contribution of each category was calculated using the Monte Carlo approach described above for enrichment with the following modifications:

6. In step 3: for each gene, the number of neutral mutations was also simulated from a Poisson distribution with rate parameter equal to the gamma-simulated rate parameter multiplied by the probability of a mutation occurring in the gene. Conceptually, this

mutation count is simulated under the hypothesis of *neutral selection* on the mutations within the gene.

- In step 4: the excess for each gene is calculated as the difference between the number of mutations simulated under positive selection and the number simulated under neutral selection. The total excess for each mutation category is summed across all genes and the relative contribution calculated as above.

*Analysis of alternative splicing events in RNA-seq data*

We obtained RNA-seq data for 9 samples carrying deep intronic predicted cryptic splice SNVs (i.e., distance to nearest exon boundary > 20 base-pairs) in TSGs with a significant burden of predicted noncoding cryptic splice SNVs. This represented all such carriers with available RNA-seq data. We downloaded the STAR aligned BAM files for each donor and six randomly selection non-carriers from the same cancer cohort, and we used bedtools bamtofastq to convert these reads into FASTQ files for de novo alignment. We then ran olego[27] with the default junction database and max edit distance of 4 (flag -M 4) on each FASTQ file. Olego is specifically designed for increased sensitivity to de novo splicing in RNA-seq reads. The de novo aligned sam files were then converted to bam files, sorted, indexed, and processed for junctions by Regtools[28] for downstream analysis (input parameters: -a 8 -m 50 -M 50000). For each of the carrier-control pairs, we performed differential splicing analysis using LeafCutter as described by Li et al.[29]. The introns in each pair were clustered using the leafcutter_cluster_regtools.py script, requiring a single split read to support a junction and assuming a maximum intron length of 500Kb (input flags -m 1 -o -l 500000). Differential splicing was then evaluated using the leafcutter_ds.R script using the Gencode v19 exons provided with the software. When a gene had more than one transcript available, we used the canonical transcript as annotated in UCSC genome browser. We considered a predicted splice SNV to have strong supporting evidence if LeafCutter reported a splice cluster containing the predicted splice SNV that had significantly different usage between carrier and control ($p<0.05$) in the majority of the carrier-control pairs. If LeafCutter did not report a cluster containing the predicted splice SNV, we additionally examined the raw junction files from Regtools. We considered a predicted SNV to have some supporting evidence if junctions supporting the prediction were observed in the raw junction files. Two of the nine samples were discarded due to insufficient coverage of the gene of interest.

#### Noncoding driver mutations in 5' UTRs

*Discovery of promoters with a burden of indels*

We assessed the burden of indels within the set of promoters defined by the PCAWG consortium[1] using Dig with default parameters. We tested the PCAWG pan-cancer cohort, excluded samples with >3000 coding mutations from this analysis.

*Discovery of multi-exonic 5' UTRs with burden of mutations*

We examined 5' UTRs that spanned multiple exons of the canonical transcripts of genes (as defined by UCSC genome browser for GRCh37) for a burden of SNVs or indels using Dig run with default settings. We additionally included the splice regions of the 5' UTRs in our analysis, defined as the 20 base-pairs bordering the start or end of an exon. We tested the PCAWG pan-cancer cohort, excluding samples with >3000 coding mutations from this analysis. To boost statistical power by limiting the number of hypotheses tested, we performed two restricted analyses considering n=106 tumor suppressor genes and n=95 oncogenes in the CGC with multi-exonic 5' UTRs.

*Analysis of mutations from Hartwig Medical Foundation cohort*

We downloaded somatic mutations observed in the Hartwig Medical Foundation metastasis cohort[30] from their online data portal (https://database.hartwigmedicalfoundation.nl/), excluding skin and hematopoietic tumors. Since we could only download mutations specific to a gene, we did not quantify burden with Dig. Rather, we directly compared the rate of SNVs in the 5' UTR, first intron, and 1kb upstream region of *ELF3* to the rate of synonymous mutations in *ELF3* using a Fisher's exact test.

*Analysis of expression levels*

We obtained gene expression levels (FPKM) and gene-level copy number estimates from the PCAWG data portal for all tumors for which RNA sequencing was performed. For a gene of interest, we applied a fixed-effects linear regression model to residualize the expression values for gene-level copy number per sample and the interaction between gene-level copy number and the cancer project that originally generated the RNA-seq data. We then normalized the residual expression values to have mean zero and unit variance across all samples and compared the normalized values between mutation carriers and noncarriers using a Mann-Whitney U-test.

*Analysis of alternative splicing*

Alternative splicing in carriers of *TP53* exon 1 5' UTR splice region mutations for which RNA-seq data existed was evaluated as described for carriers of noncoding cryptic splice mutations.

#### Driver mutations in long-tail cancer driver genes in whole-exome and targeted sequenced cohorts

We estimated the neutral mutation rate of activating SNVs in 69 oncogenes and pLoF SNVs (nonsense and canonical splice) in 56 TSGs that were common to all whole-exome and targeted sequencing samples in each cancer cohort that we had assembled (see above).

Because synonymous mutations were not available from the targeted sequenced samples, we instead used missense mutations with CADD phred score <15 to estimate the scaling factor. Specifically, for a give cancer type we calculated the number of neutral mutations with CADD phred <15 expected under mutation map for each cancer across 51 TSGs common to all whole-exome and targeted sequenced samples (and excluding five genes with a burden of CADD phred<15 missense SNVs in whole-exome sequenced samples). We then counted the number of observed missense mutations with CADD phred <15 in those genes in the target cohort and estimated the scaling factor as the number observed divided by the number expected. Considering only whole-exome sequenced samples (for which synonymous SNVs were available), we showed that scaling factor estimation by CADD phred <15 missense SNVs in targeted sequenced TSGs yielded similar results to estimation of the scaling factor by synonymous mutations across all genes excluding *TP53* (**fig. S17**, **table S22**). As expected, TSGs did not carry an excess of CADD phred<15 SNVs in our megacohorts under this scaling factor (fig. S17, table S23). Moreover, long-tail oncogenes did not have an excess of pLoF mutations in any of the cancer cohorts under this scaling factor, consistent with the expectation that pLoF mutations in oncogenes are neutral (**fig. S17**, **table S24**).

For each type of cancer, we assembled a list of known driver genes identified in any of three recent, pan-cancer driver gene discovery efforts[3,31,32]; we considered only genes that were discovered with FDR significance <0.1, the significance threshold common across the driver element detection literature. For a given cancer, we considered any gene not included in the list of known driver genes for that cancer type to be a “long-tail” gene. We directly estimated the P-value of the mutational burden long-tail genes by convolving the neutral mutation distributions for each individual gene and calculating the upper-tail probability of at least the number of observed mutations across all genes occurring by chance under the null distribution. We calculated 95% confidence intervals of excess mutations using the same Monte Carlo approach as for calculating the 95% confidence intervals of excess genic mutations in our analysis of cryptic splice SNVs. We divided the estimated excess per cancer type by the total number of samples in the cohort to estimate an excess rate per sample.

#### Power analysis

We conservatively simulated Dig's power to detect driver SNVs at different carrier frequencies across particular genomic elements (lncRNAs from PCAWG, cohesion loop boundaries from ENCODE[33], noncoding cryptic splice sites, and activating SNVs) under the pan-cancer mutation model using the following Monte Carlo approach:

For a given target sample size and carrier frequency:

1. For each element, randomly draw a mutation rate parameter from the gamma distribution defined by mean and variance estimated by the kilobase-scale model.
2. For each element, estimate the scaling factor as the target sample size divided by the pan-cancer sample size (n=2279) and randomly draw an observed number of mutations from a Poisson distribution with rate parameter equal to the sampled rate multiplied by the scaling factor and by the probability of an SNV in the element.
3. For each element, randomly sample the number of driver mutations from a Poisson distribution with rate parameter equal to the target sample size multiplied by the carrier frequency.
4. Count the number of elements for which the sum of the background mutations and driver mutations exceeded the Bonferroni-corrected $\alpha$ <0.05 threshold under Dig's negative binomial null mutation distribution for each element. Divide the count by the total number of tested elements to estimate a detection likelihood.
5. Repeat steps 1-4 one thousand times and average the detection likelihoods across all simulations.

### Supplementary notes

#### 1. Comparison of cancer driver detection methods

Because our approach identifies driver candidates by testing for selection, we compared its accuracy to other methods that also use test for selection. We first compared our method's ability to identify driver genes in the PCAWG dataset against MutSigCV[5] and dNdScv[2], two widely used methods created specifically to identify genes under positive selection. Following previous works[3,19], we used the Cancer Gene Census (CGC)[21] as a conservative approximation of the true-positive rate and found our method matched or exceeded the F1-score (a joint measure of sensitivity and specificity) of the other methods in 24 of 32 PCWAG cohorts (excluding hematological and skin malignancies[19]) (uniquely highest score in 13 cohorts; tied for highest in 11 cohorts) (**fig. S7**, **table S7**). We additionally calculated the receiver-operator curves for the top 600 genes identified by each method in the PCAWG pan-cancer cohort and found Dig systematically identified more true-positive drivers and fewer false-positives than the other methods (**fig. S7**), a pattern that we also observed when we additionally compared the methods across whole-exome sequenced (WES) cohorts[3] (**fig. S8**). We additionally found that Dig's ability to accurately recall noncoding drivers previously identified in the PCAWG dataset was comparable to that of three other burden-based non-coding driver detection methods, Larva[20], ActiveDriverWGS[22], and DriverPower[19] (**fig. S9**, **table S8**), although this analysis was biased against Dig because the other three methods were used to generate PCAWG's own set of noncoding drivers.

#### 2. Variance estimation in deep neural networks with Gaussian processes

While it is intuitive that more accurate prediction of the expected neutral mutation rate can improve power to identify drivers, the accuracy of variance prediction also plays a crucial role, particularly in ensuring well-calibrated p-values. We previously investigated the accuracy of the

CNN+GP architecture to estimate kilobase-scale mutation rates compared to other architectures in a simulated dataset (full details in Yaari et al.[18]). Here we review the results about variance because they provide additional insight into reasons underlying our methods power to detect driver events.

In brief, we generated a synthetic kilobase-scale mutation rate dataset for the PCAWG melanoma, esophageal, and stomach cancer datasets using a k-nearest-neighbors strategy. For each 50kb window, we found its 500 nearest neighbors based on mean epigenetic context from the Roadmap Epigenomics dataset and defined the "true" mean and variance for that window to be the mean and variance of the mutation rate across the 500 nearest neighbors. We then trained methods to predict the mean and variance of each window based on a mutation count randomly simulated from a negative binomial distribution defined by that true mean and variance. We then compared the predicted mean and variance to the true mean and variance for each method (**fig. S23**).

The simulated dataset has an interesting feature: the variance of the mutation rate plateaus beyond a certain expected mutation rate. That is, while the expected mutation rate continues to increase, the variance of the mutation rate does not increase. An important feature of a Gaussian process is that it *nonlinearly* predicts mean and variance; the relationship between mean and variance is not imposed a priori. This has the effect of enabling the CNN+GP estimation method to learn that variance plateaus as expected mutation rate increase (**Fig. S23c**). Thus, statistical tests remain well powered to identify outlier events even at high mutation rates. This is not the case for the linear regression methods often employed to predict the mean and variance of mutation rate. For example, negative binomial regression imposes a quadratic relationship between mean and variance: $\sigma^2 = \mu(1 + \beta \cdot \mu)$, where $\beta > 0$ is the estimated overdispersion parameter. This has the effect of forcing the variance to increase *faster than* the expected mutation rate, leading to variance estimates considerably larger than the true variance when expected mutation rate is high (**Fig. S23c**). Thus, negative binomial regression loses power to detect outliers in high mutation rate contexts.

#### 3. Additional details on alternative splicing analysis with LeafCutter

Of the nine predicted cryptic splice SNV carriers for which we obtained RNA-seq data (**Methods**), two carriers were discarded due to insufficient coverage either at the gene of interest (DO222305, median coverage of *CIITA* of 17 reads) or globally (DO9074, median depth of coverage of 33). Of the remaining 7 carriers, 4 had clear evidence of alternative splicing: LeafCutter[29] reported a splicing cluster containing the predicted splice SNV with significantly different usage ($P < 0.05$) between the carrier and at least a majority (4 of 6) of the control pairs (**table S12**). We further investigated the remaining 3 predicted cryptic splice SNV carriers and observed that 2 of the 3 had some evidence of alternative splicing. For one of these carriers (DO44111), LeafCutter reported significant alternative splicing in the carrier compared to half of the controls; however, it was unclear if the difference was specifically related to the predicted splice SNV. The other carrier (DO52675) had evidence of differential splicing that was not reported by LeafCutter. Specifically, by manually annotating the junction files produced by Regtools[28] with the introns defined in ENSEMBL, we observed that the carrier used an alternative site consistent with the predicted splice SNV in approximately 10% of transcripts, while the controls utilized this site in approximately 1% of transcripts. The remaining (DO33392) sample did not have evidence of alternative splicing upon manual review.

#### 4. Investigation of mutational burden in *ELF3* 5’ UTR

The PCAWG consortium previously carefully reviewed noncoding mutational hotspots in the PCAWG dataset[1] and cataloged several reasons for excess mutations that were unrelated to positive selection: activation-induced cytidine deaminase (AID) activity in lymphomas, impaired nucleotide excision repair (NER) at transcription factor binding sites in melanomas, activity of endogenous apolipoprotein B mRNA- editing enzyme catalytic subunit (APOBEC) family deaminases, particularly in the in the loop region of predicted hairpin structures, and systematic short-read mapping inaccuracies leading to artefactual mutation calls. We examined whether any of these processes could be responsible for the observed enrichment of SNVs in the 5'UTR of *ELF3*.

In our analysis of the 5' UTR of *ELF3*, we specifically excluded hematopoietic tumors and melanomas, so neither AID nor NER likely account for the observed elevated mutation rate. To investigate the possible role of APOBEC at the 5'UTR of *ELF3*, we obtained the results of the ABOPEC analysis performed by the PCAWG consortium in which each observed mutation was annotated for whether it could be attributed to APOBEC. Of the six SNVs observed in the *ELF3* 5' UTR, only one was annotated as occurring in a context targeted by APOBEC; however, the sample in which that mutation occurred was not significantly enriched for APOBEC mutations of that kind nor did the mutation occur within a cluster as would be expected if it were due to APOBEC mutagenesis. We thus do not believe APOBEC likely explains the mutational excess in the *EFL3* 5' UTR. We next examined the gnomAD database[34] which both cataloged population polymorphic germline genetic variation and noted regions of the genome where mapping artefacts were present. The 5' UTR of *ELF3* was not annotated as a region with mapping artefacts by gnomAD. Moreover, of the 16 somatic mutations observed in the PCAWG and Hartwig datasets, only one affected a position also affected by a germline SNP (the canonical splice site chr1:201979836, although the mutation itself is different). The germline SNP was rare (2 alleles observed in >30000 haplotypes). Moreover, the six mutations in the PCAWG dataset were observed in five different cancer types and the ten mutations in the Hartwig dataset were observed in seven different cancer types. Thus, the enrichment cannot be attributed to a mutational process specific to one cancer type. Finally, the mutation enrichment was specific to the canonical 5' UTR of *ELF3*; enrichment was not observed in surrounding regions as was noted by PCAWG for several lncRNAs. In summary, we were unable to explain the mutation burden observed in the 5' UTR of *ELF3* by any processes that had been previously noted to increase mutation rate independent of positive selection.

#### 5. Preliminary analysis of enhancer networks

An analysis of the SNV and indel burden in enhancers (obtained from Nasser et al.[35]) of 725 CGC genes using Dig with default settings revealed 36 enhancers with significant (FDR<0.1) mutational burdens. To coarsely filter regions potentially affected by unmodeled local hypermutation processes, we required that observed mutations each occur in a unique sample. This filter reduced the number of enhancers to ten (**table S20**). Two enhancers (for *LEPROTL1* and *SRGAP3*) contained recurrent mutations (*LEPROTL1*: 8:29952919-G>A (n=7), 8:29952921-C>A,G,T (n=5); *SRGAP3*: 3: 8486222-G>C,T (n=6)); however, it is possible that these mutational hotspots could result from *APOBEC* mutagenesis or mapping artefact[1]. Carriers of mutations in several enhancers demonstrated significant (P<0.05) or nearly-significant (P<0.1) differences in expression compared to non-carriers (not corrected for multiple hypothesis testing). For example, carriers of mutations in the *NCOR2* enhancer (12:125422682-125425761) had a nearly significant decrease in expression (P=0.078). However, expression did not always change in a direction consistent with the known or predicted function of the gene in tumorigenesis. For example, carriers of indels in the *MSI2* enhancer (17:54992281-54993673) had decreased *MSI2* expression (P=0.0081) based on carrier tumors from kidney, rectum, and ovary; however, *MSI2* is a known oncogene in hematopoietic cancers. More follow-up analysis will be necessary to determine whether the mutational enrichment constitutes positive selection or unaccounted for neutral mutational processes.

### Supplementary Figures

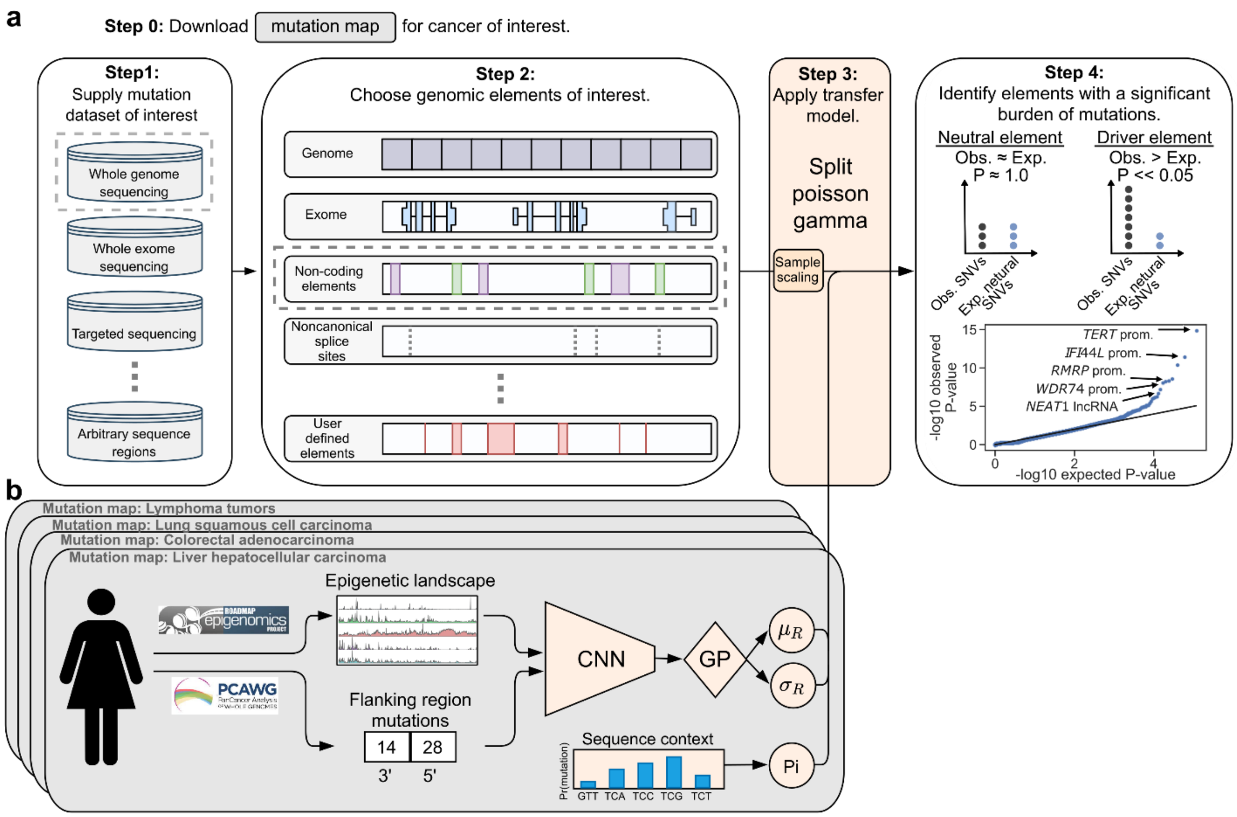

**Supplementary figure 1: Detailed overview of the Dig model. a,** Dig takes as input somatic mutations (SNVs and/or indels) (**Step 1**) identified from a cancer cohort sequenced with any methodology and a set of genomic elements of the user's interest (**Step 2**). The neutral mutation rate from an available neutral somatic mutation map (detailed in panel **b**) is transferred to the selected SNV dataset via a closed-form probabilistic model that infers only a single scaling parameter at runtime (**Step 3**); then, a P-value for positive selection is calculated for each element by comparing the number of observed mutations to the number of expected neutral mutations (**Step 4**). **b,** A neutral mutation map for a particular cancer consists of 1) the mean and variance of the number of neutral mutations in kilobase-scale regions of the genome (default: 10kb) as inferred by a convolutional neural network (CNN) and Gaussian process (GP) based on 735 epigenetic features from the Roadmap Epigenomics dataset and ENCODE (and optionally the number of mutations observed 100kb up- and downstream of the region in the a training cancer cohort dataset); and 2) a sequence context model that provides the genome-wide likelihood of any mutation given its sequence context (default: trinucleotide sequences).

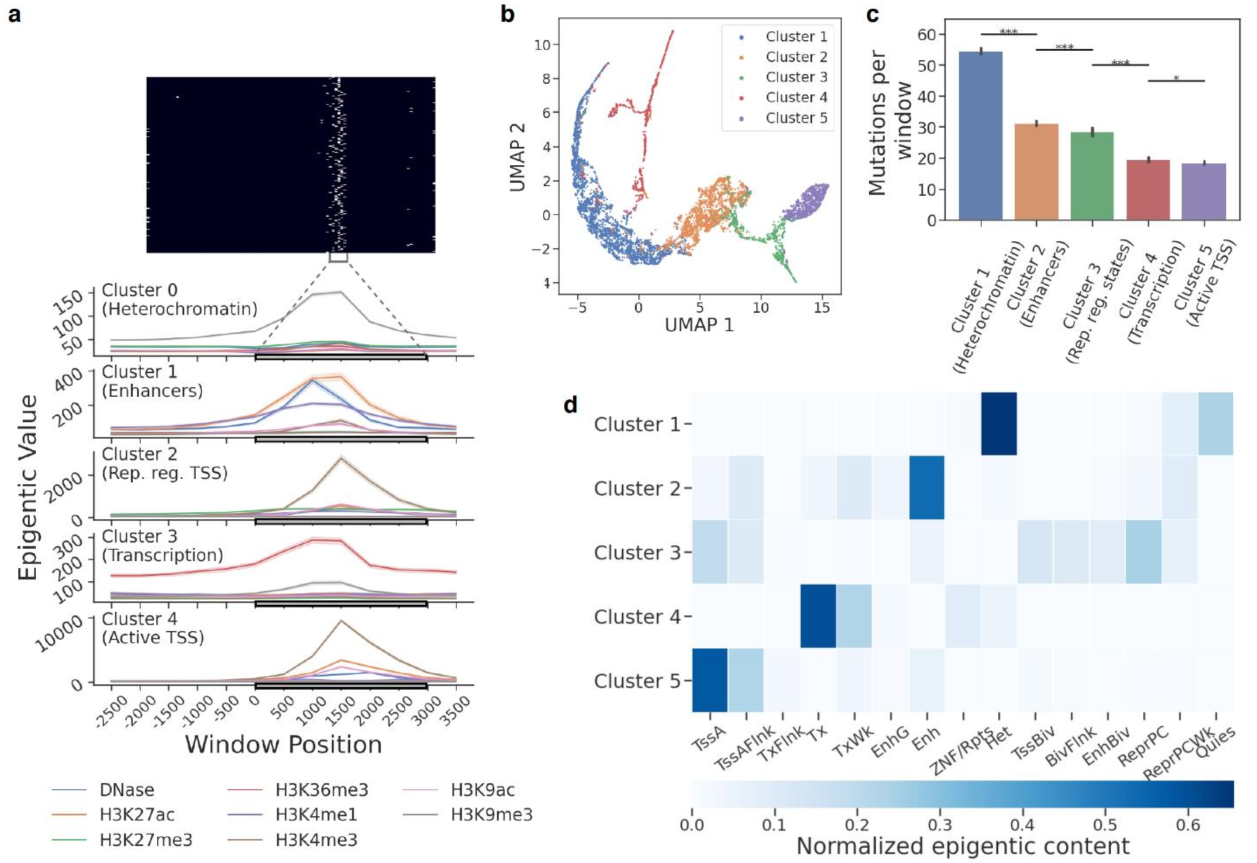

**Supplementary figure 2: Epigenetic input features used by Dig to predicted mutation density in lung squamous cell carcinoma**. **a,** an example of a feature map with an attention column produced by Dig and the mean chromatin content of five epigenetic features consistently identified by Dig as salient to prediction accuracy. Grey rectangles denote the 95% confidence interval of the width of the attention column across all regions. **b,** UMAP visualization and clustering of the input space attended to by dig across 50kb regions. **c,** The mean number of mutations in regions containing an attention column from a given cluster. Error bars: 95% CI. **d,** The average epigenetic content of each cluster as determined by epilogos[24].

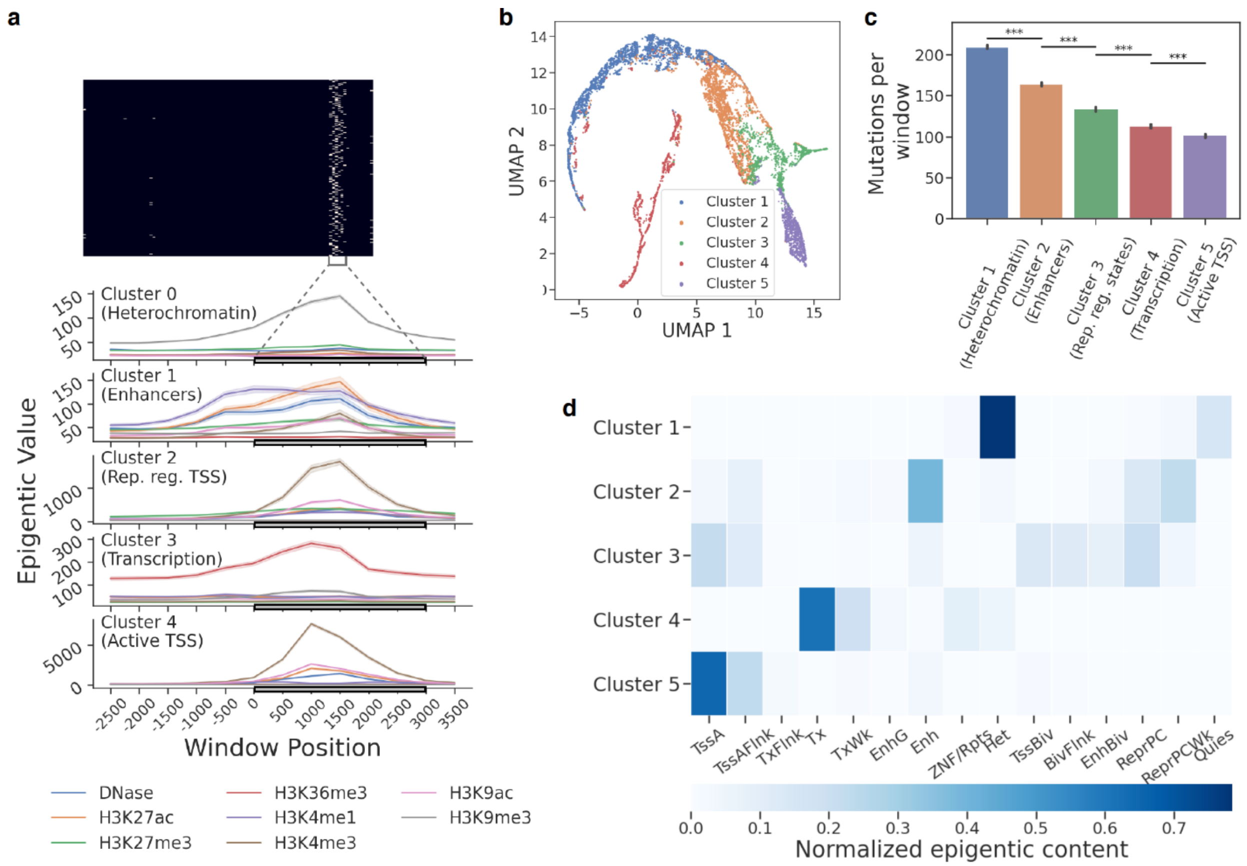

**Supplementary figure 3**: as in Supplementary Fig. 2 but for colorectal adenocarcinoma (excluding MSI-high samples).

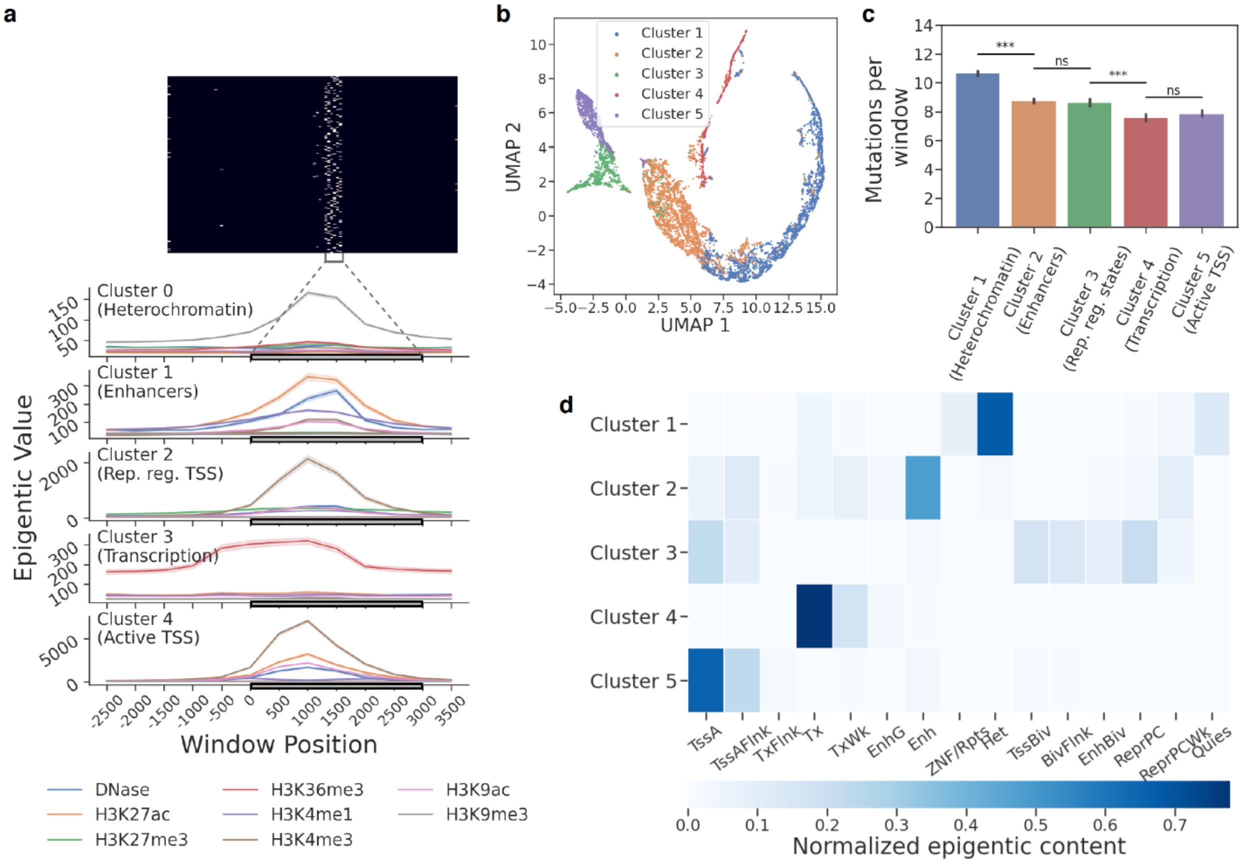

**Supplementary figure 4**: as in Supplementary Fig. 2 but for bladder adenocarcinoma

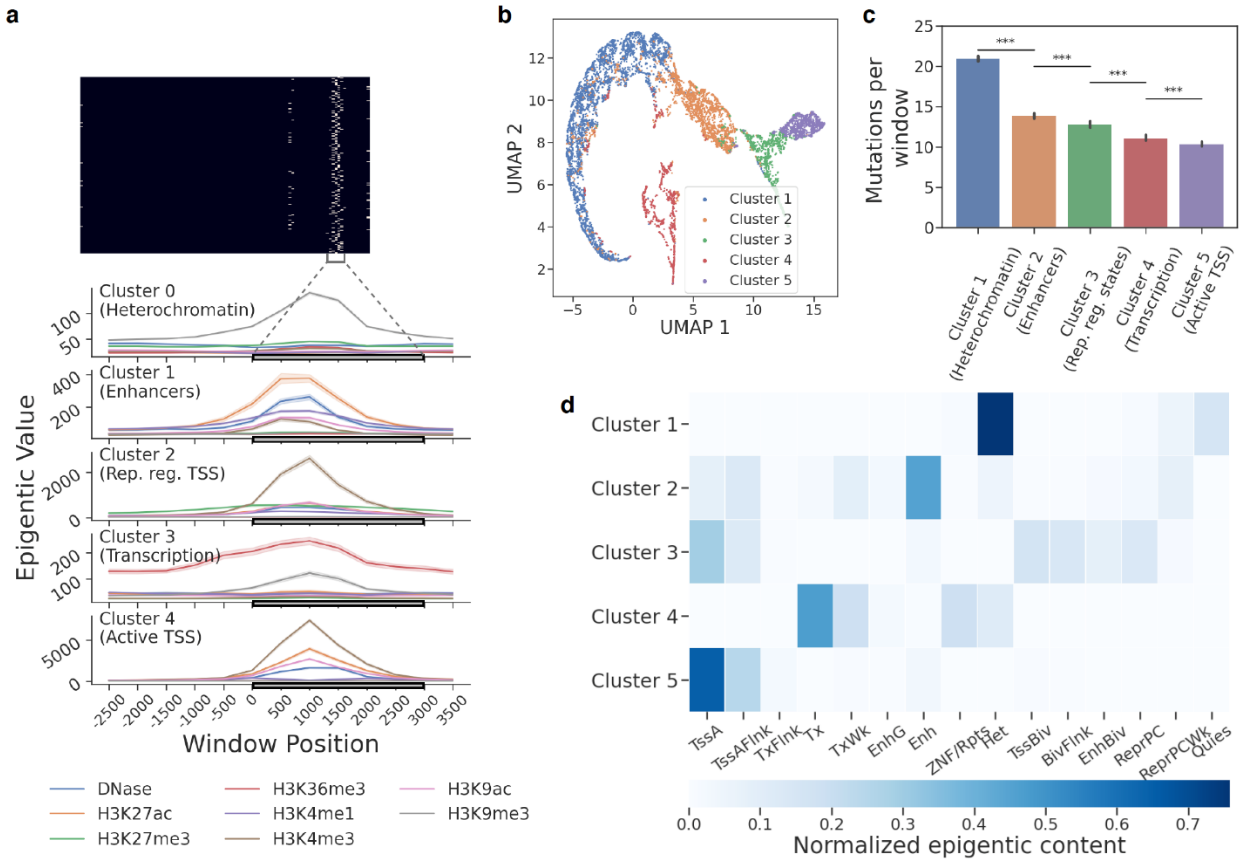

**Supplementary figure 5**: as in Supplementary Fig. 2 but for head and neck squamous cell carcinoma.

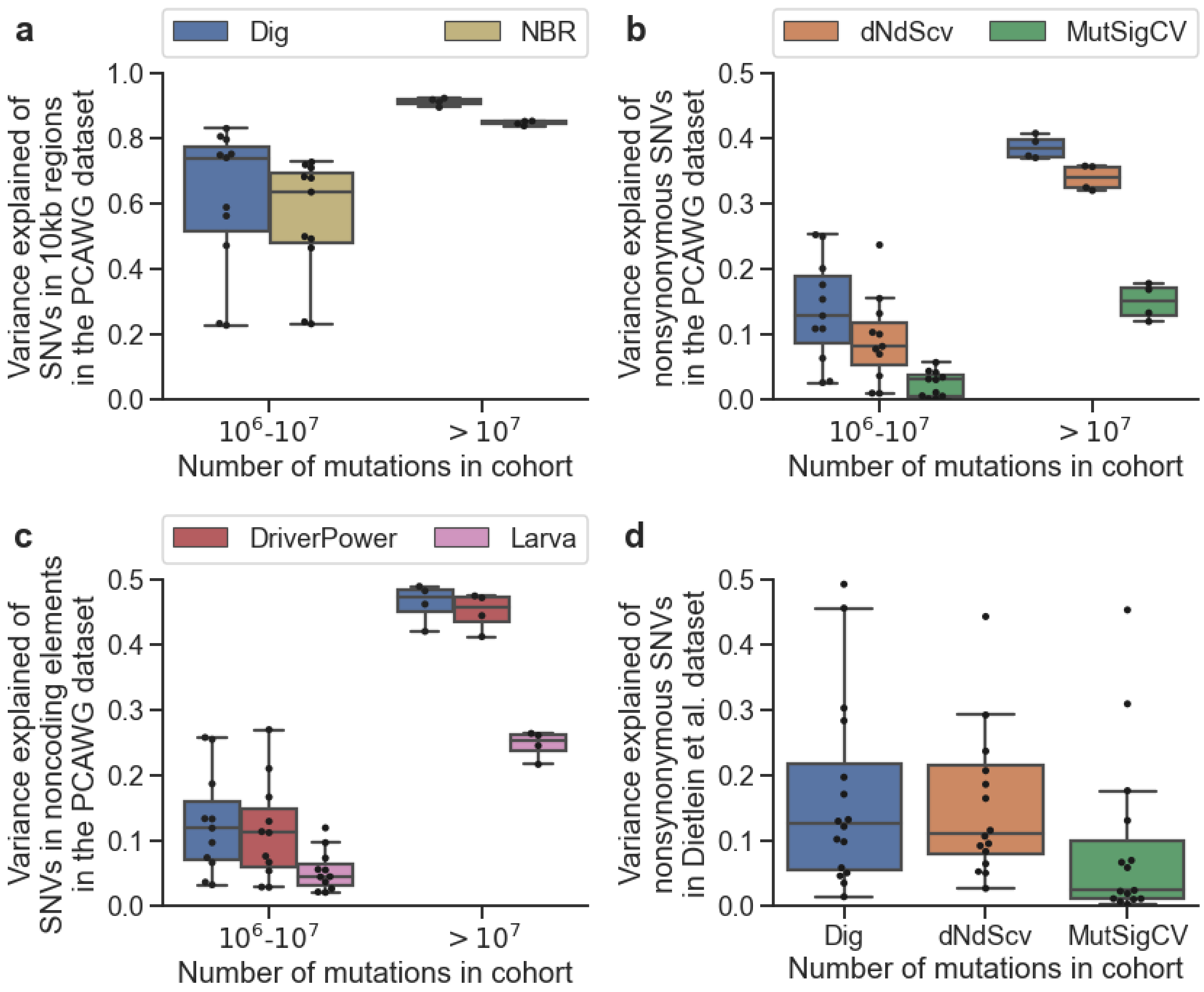

**Supplementary figure 6: Comparison of variance explained of SNV counts across methods, annotations, and cohorts. a,** Variance explained of SNV count in 10kb regions tiled across the genome by Dig and NBR[1] in 16 PCAWG cancer cohorts with >1 million SNVs (excluding hemopoietic tumors, for which NBR failed for converge). Regions in which <50% of 36mers are unique are excluded as are regions in the 99.99$^{th}$ percentile of mutation count. **b,** Variance explained of nonsynonymous SNV count in genes 1-1.5kb in length (n=3,740 genes) in 16 PCAWG cancer cohorts. **c,** Variance explained of SNV count in enhancers and noncoding RNAs (long and short) 0.5-1kb in length (n=7,412 noncoding elements) in 16 PCAWG cancer cohorts. **d,** as **b** for 16 whole-exome sequenced cancer cohorts from Dietlein et al.[3].

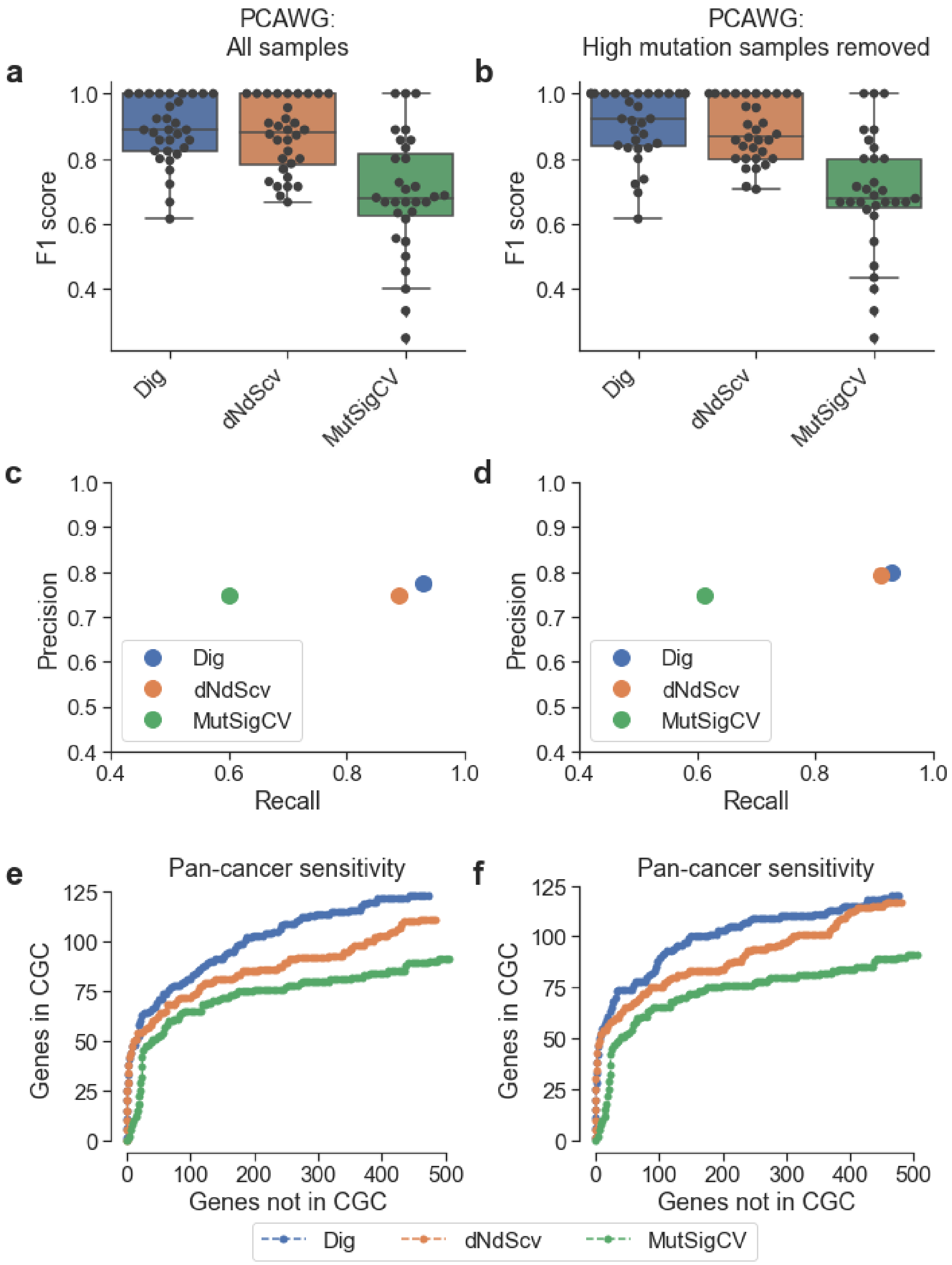

**Supplementary figure 7: Precision-recall comparison of gene driver methods in the PCAWG cohort. a,b** F1-score (harmonic mean of precision and recall) in 32 PCAWG cohorts (melanoma and hematopoietic tumors were excluded as in previous work[19]) across 16,794 genes common to the three methods. Precision and recall were calculated using genes in the Cancer Gene Census as a conservative true positive set. **a,** All samples. **b,** Excluding samples with >3000 coding mutations and restricting the total number of mutations per sample per gene to 3 (default filtering options for dNdScv). **c,d** Recall and precision measured across all 32 PCAWG cohorts for **c,** all samples and **d,** samples with <3000 coding mutations. **e,f** Receiver-operator characteristic curves for the pan-cancer cohort for **c,** all samples and **d,** samples with <3000 coding mutations.

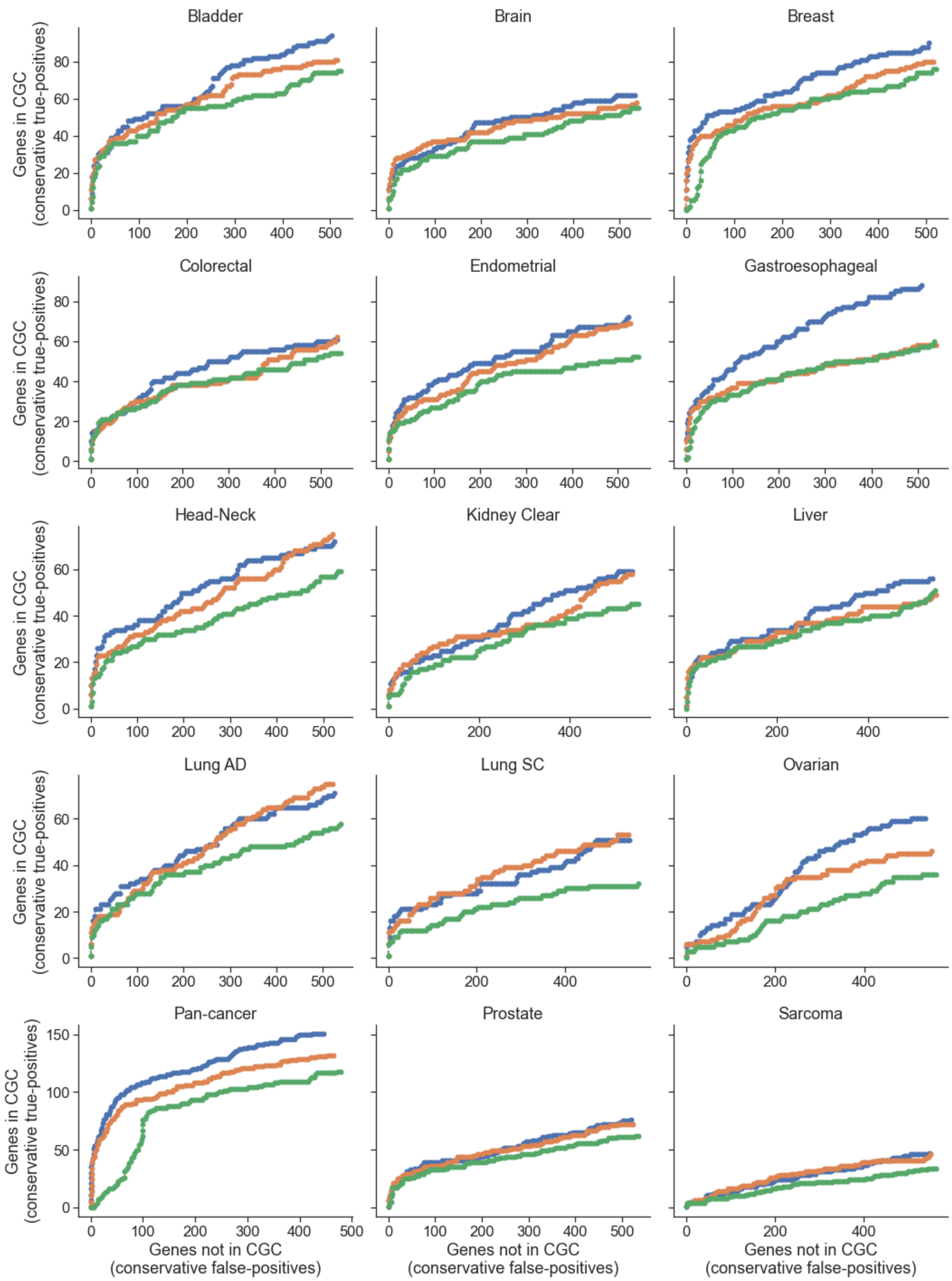

**Supplementary figure 8: Receiver-operator characteristic (ROC) curves for 15 whole-exome sequenced cohorts from Dietlein et al.**[3]. Genes in the Cancer Gene Census (CGC) were used as a conservative set of true positives. The MutSigCV model produced mis-calibrated p-values for the pan-cancer cohort, suggesting that its model assumptions may have been violated by the large cohort of heterogeneous cancer types.

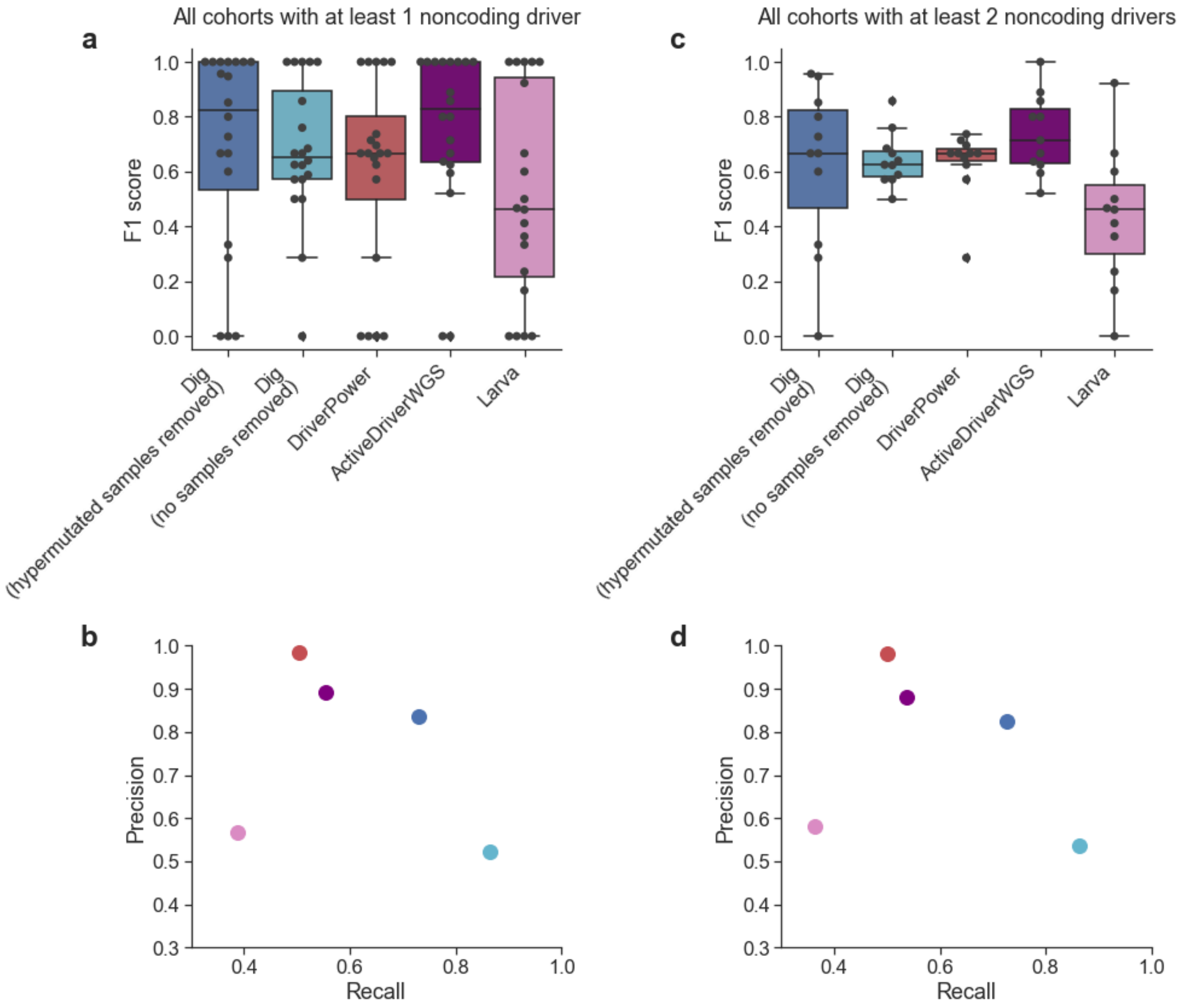

**Supplementary figure 9: Precision-recall comparison of noncoding driver detection methods in PCAWG dataset. a,** F1-score across 95,231 noncoding elements as defined in Rheinbay et al.[1] in PCAWG cancer cohorts with at least one identified noncoding driver (n=20 cohorts). The performance of Dig was also evaluated when removing samples with >1000 SNVs across all elements and restricting the total number of SNVs per sample per element to 3. DriverPower and Larva do not have built-in filtering options. ActiveDriverWGS was run with default filtering which removes any sample with >30 SNVs per megabase. **b,** Recall and precision by method combined across the cohorts in **a**. **c,d,** as in **a** and **b** but restricting to n=11 cohorts with at least two identified noncoding drivers.

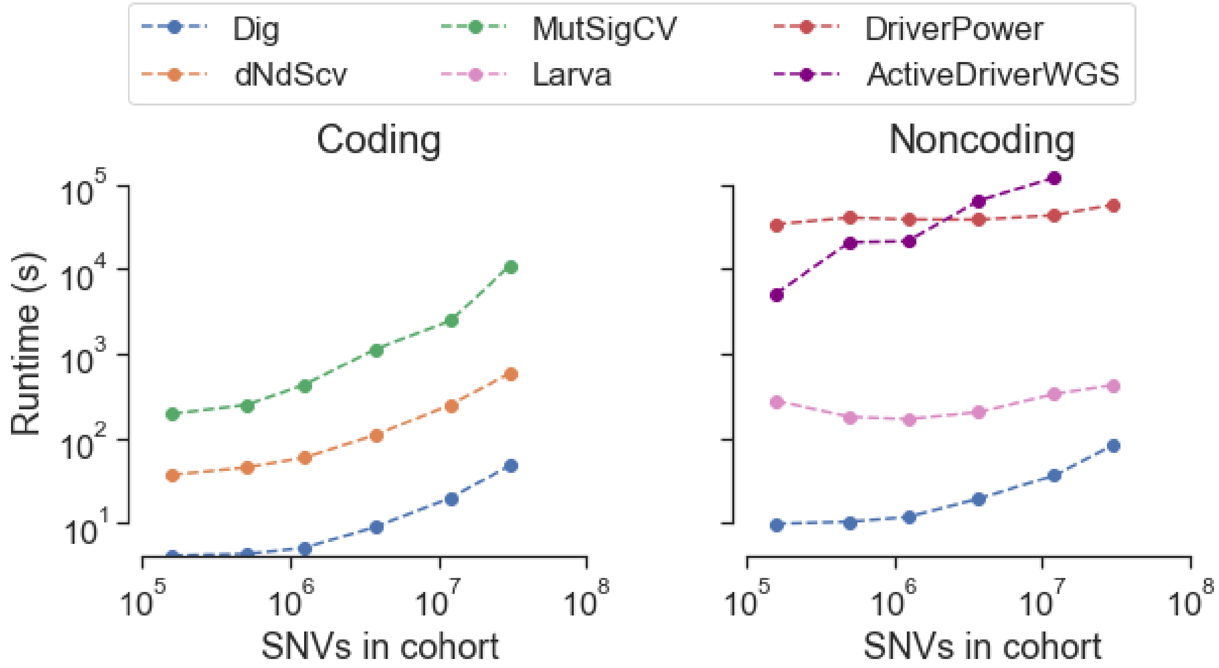

**Supplementary figure 10: Method runtime comparison**. Comparison was restricted to SNVs because not all methods analyze indels. The coding comparison was performed on N=19,210 genes for Dig and dNdScv and N=18,862 genes for MutSigCV. The noncoding comparison was performed on N=139,404 elements for Dig, DriverPower, and Larva and N=117,180 elements for ActiveDriverWGS. ActivedriverWGS required >2 days to analyze the largest cohort.

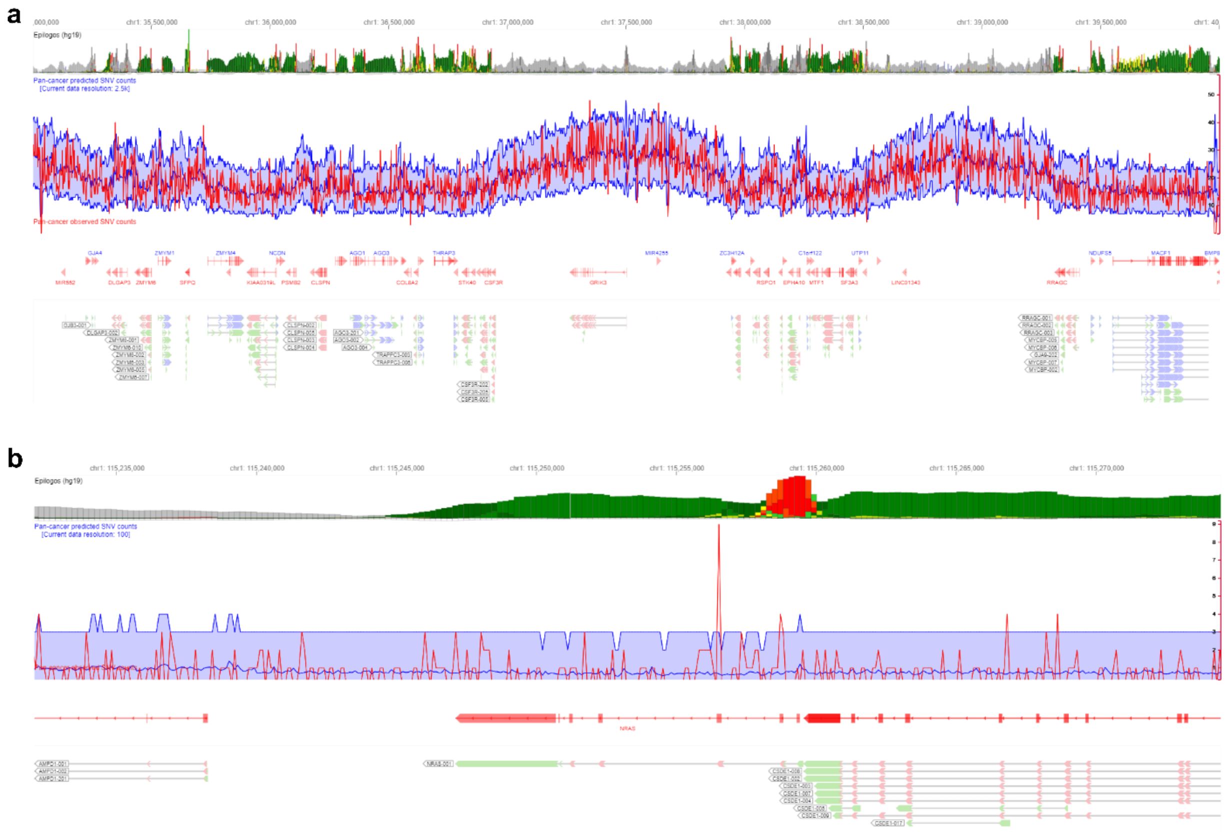

**Supplementary Figure 11: Screenshots from the online mutation map genome browser** for the PCAWG pan-cancer cohort showing **a,** a 5Mb region on chromosome 1 at 2.5kb resolution and **b,** expected and observed mutation rates around *NRAS* at 100bp resolution. Mutation enrichment in exon 3 of *NRAS* is clearly identifiable. Expected mutation rates in blue (shaded region, 95% confidence interval); observed mutation rates (from PCAWG) in red.

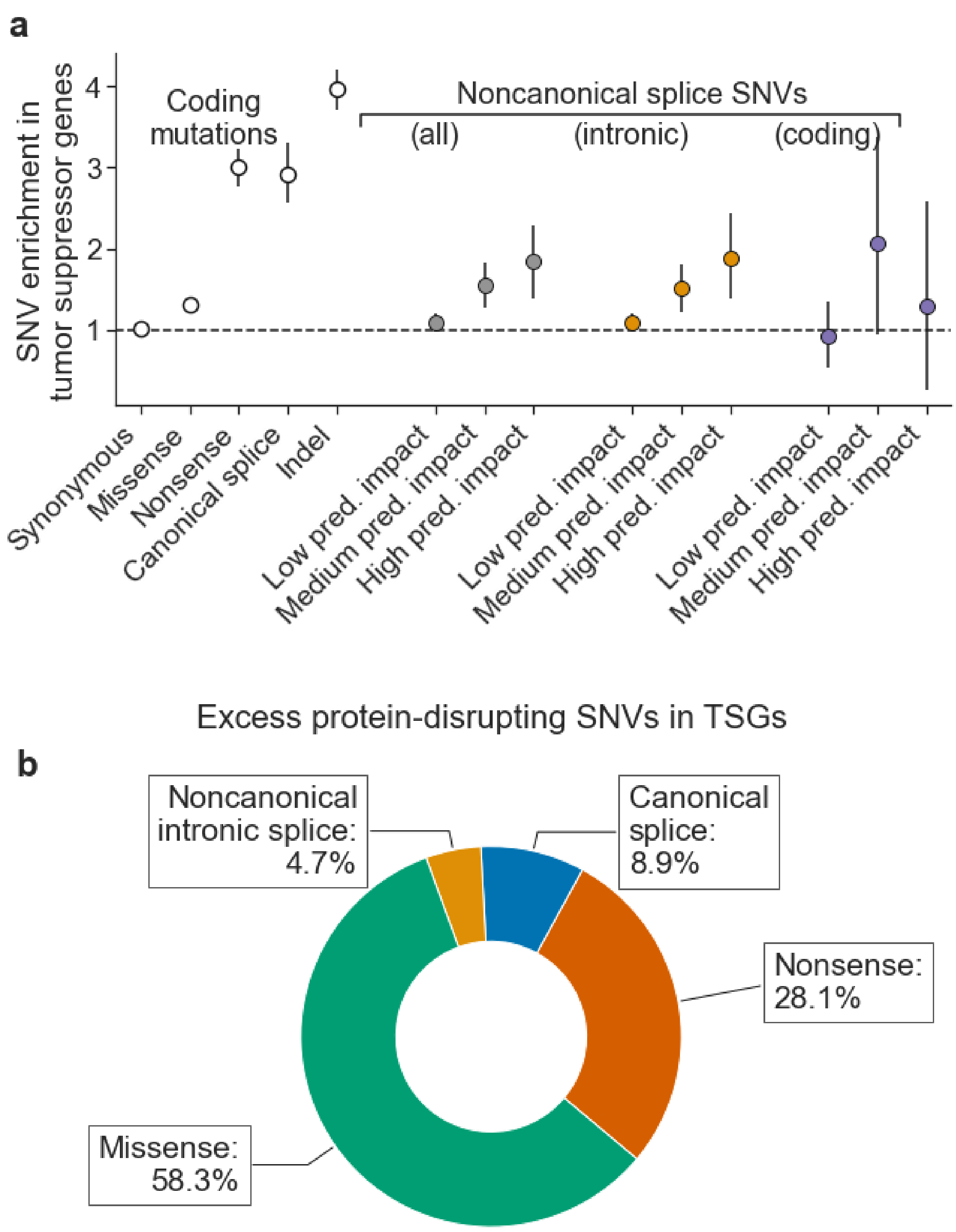

**Supplementary figure 12: SNV enrichment and excess analysis excluding samples with >3000 coding mutations. a,** as in **Fig. 3b** but excluding samples with >3000 coding mutations (default filtering criterion in dNdScv). **b,** As in **Fig. 3e** but excluding samples with >3000 coding mutations.

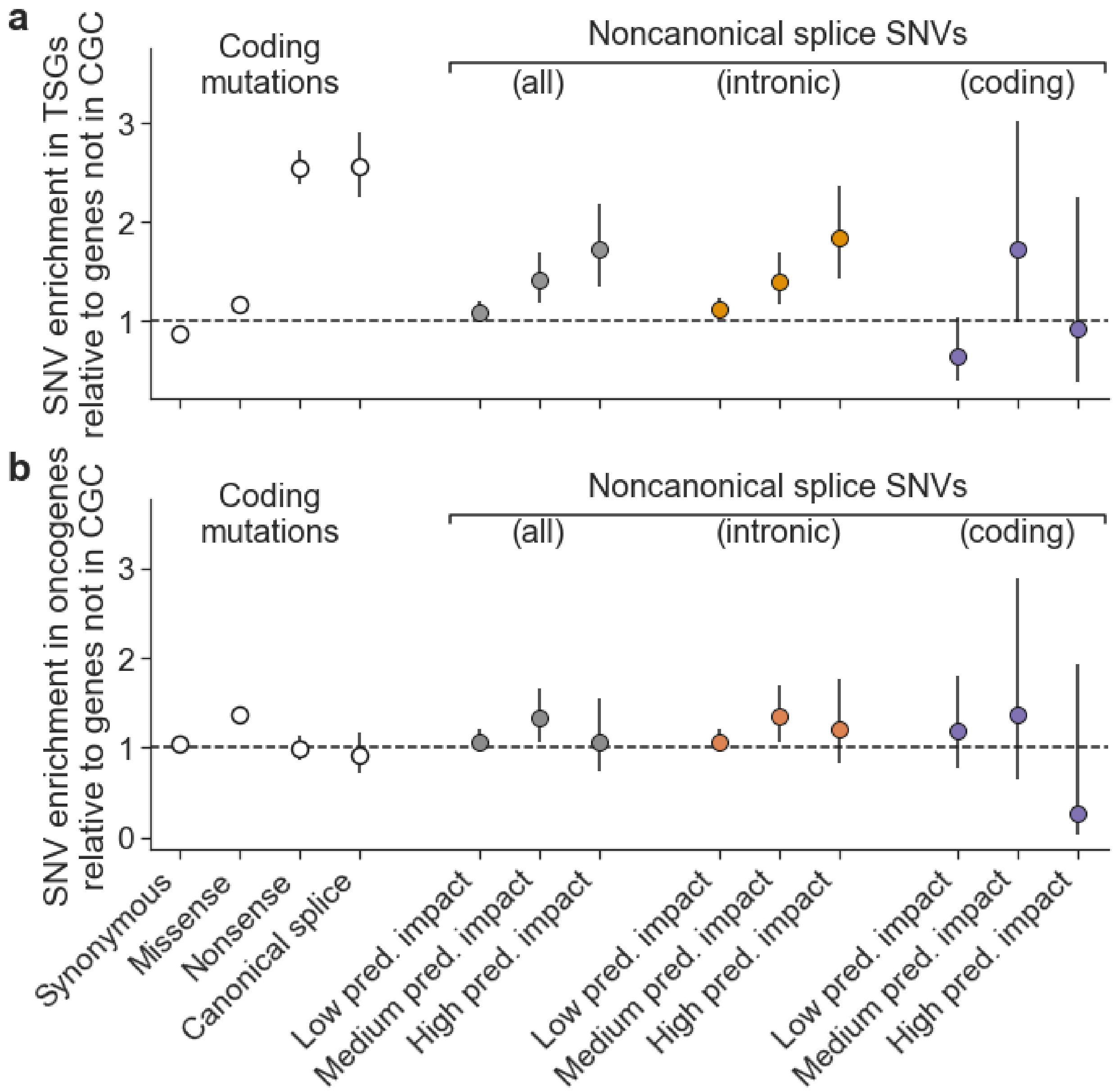

**Supplementary figure 13:** SNV enrichment analysis in tumor suppressor genes (TSGs), **a**, and oncogenes, **b**, with enrichment calculated with respect to the number of observed mutations in genes not in the Cancer Gene Census (CGC). Enrichment is calculated as the rate of SNVs of a given type observed in TSGs (oncogenes) relative to the rate of SNVs of the same type observed in genes not in the CGC.

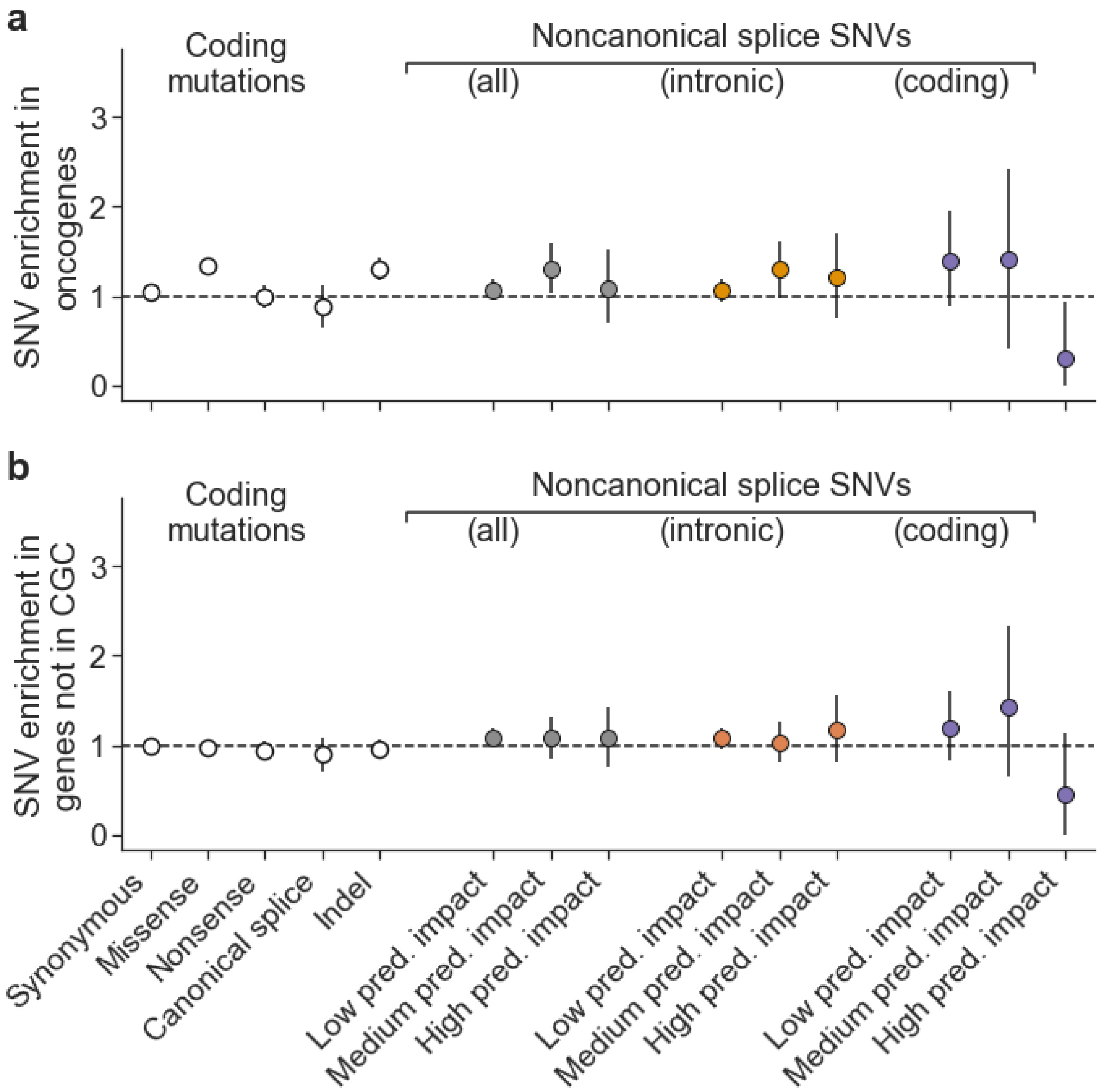

**Supplementary figure 14:** SNV enrichment analysis as in **Fig. 3b** performed for oncogenes in the CGC, **a**, and genes not in the CGC, **b**. No enrichment is significant after accounting for multiple hypothesis testing except missense mutations and indels in oncogenes, as expected.

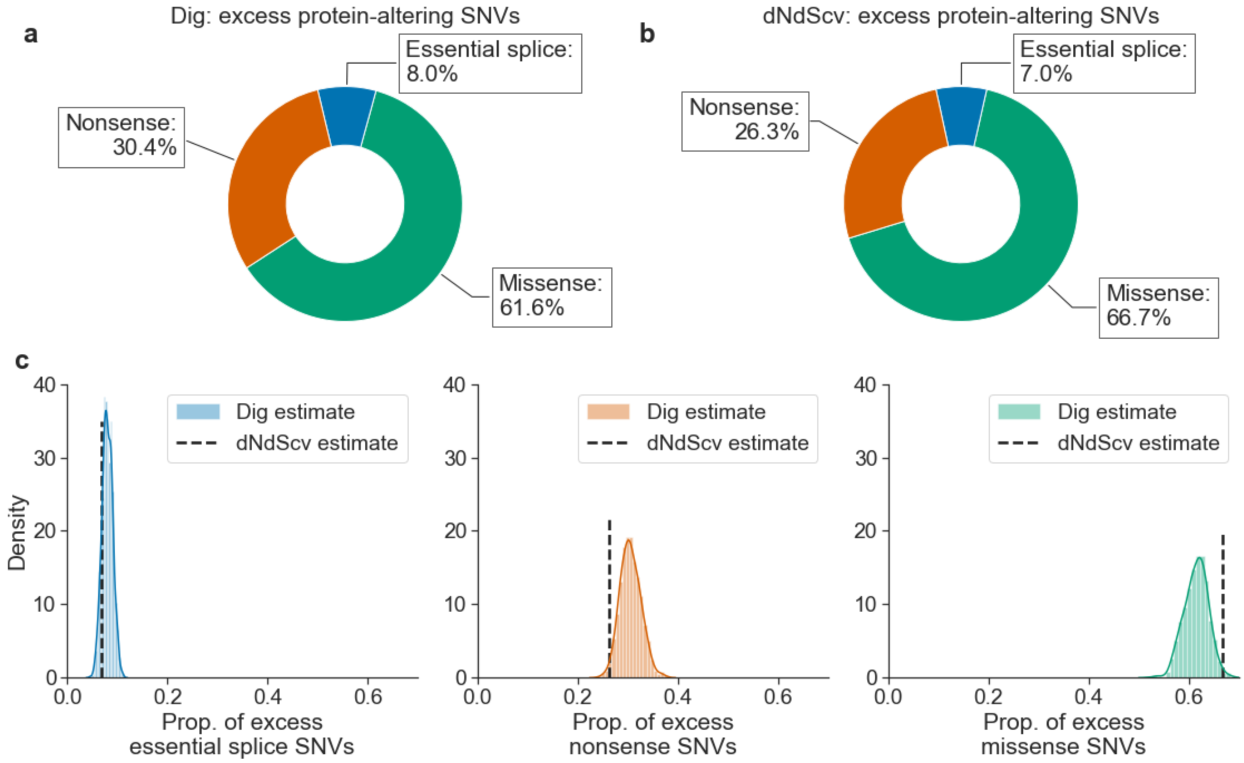

**Supplementary figure 15:** Proportion of excess protein-altering SNVs in TSGs as estimated by Dig, **a**, and dNdScv, **b**. **c,** Distribution of proportion of excess SNVs as estimated using a Monte Carlo simulation approach based on Dig (**Methods**) with the corresponding dNdScv estimate indicated with a black dashed line. Essential splice SNVs include SNVs at canonical splice sites (see **Fig. 3a**) and SNVs 5 bp 5' of an exon start, which dNdScv also considers in its analysis of splice mutations.

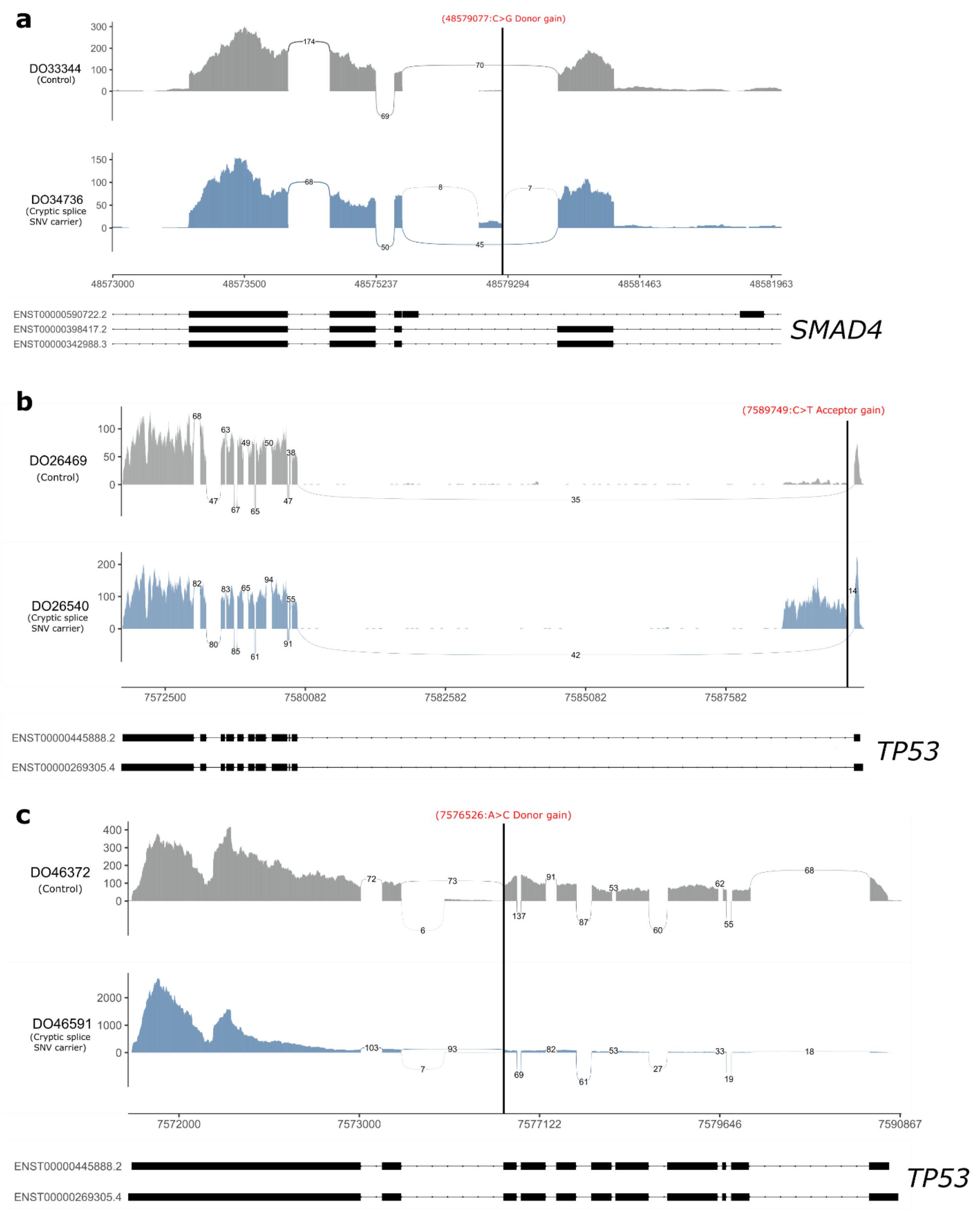

**Supplementary Figure 16: Additional predicted cryptic splice SNV carriers in which LeafCutter identified strong evidence of alternative splicing**. The location of the predicted cryptic splice SNV is marked with a thick black vertical line and labeled in red. **a,** *SAMD4* cryptic splice carrier carrier. **b,c** *TP53* cryptic splice SNV carriers.

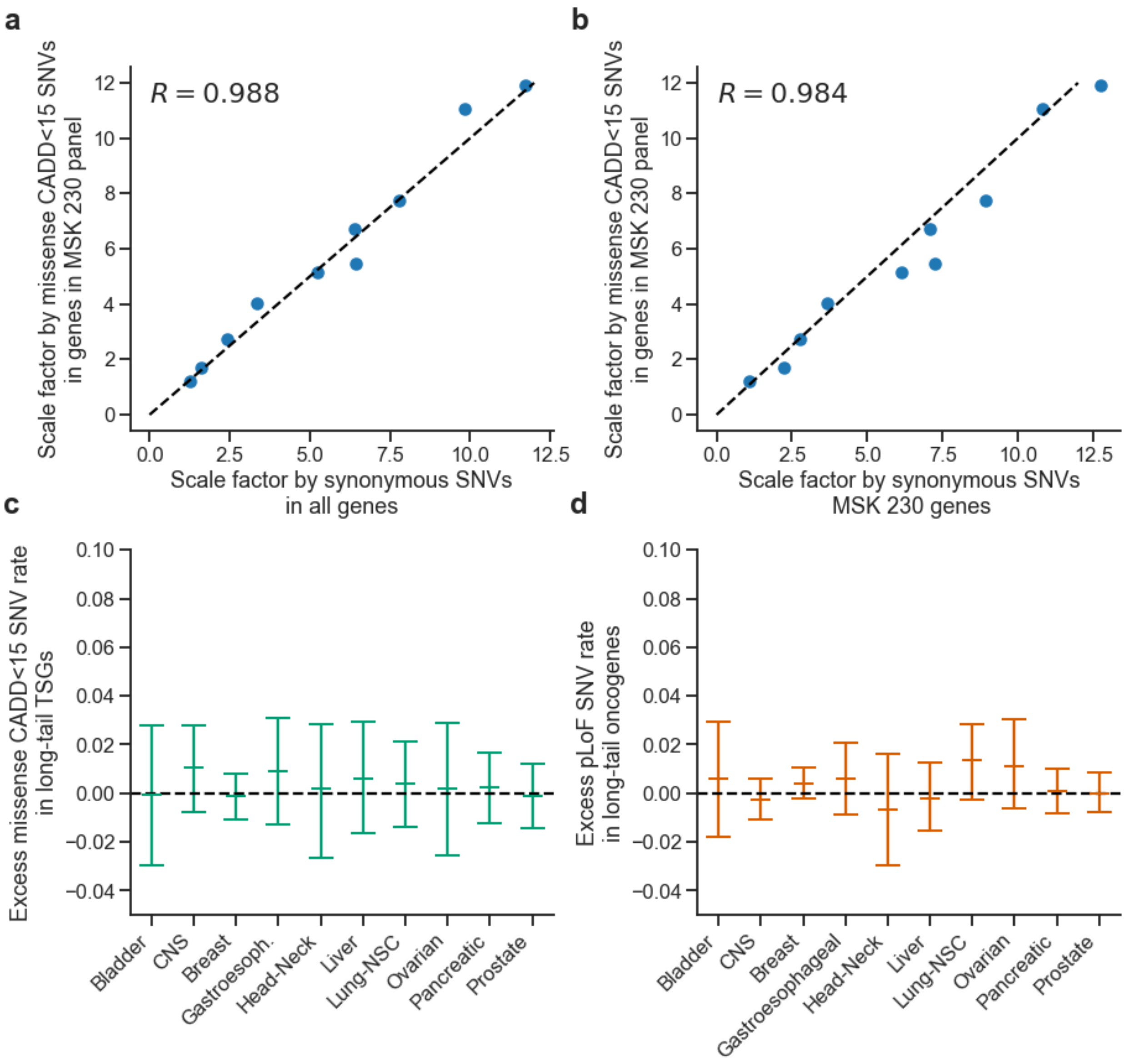

**Supplementary figure 17: Evaluation of neutral mutation model for ten solid cancer megacohorts.** Using whole-exome sequenced samples, we compared the accuracy of estimating the scaling factor based on missense SNVs with CADD phred<15 observed in genes in the MSK IMPACT 230 targeted sequencing panel (the approach used for analyzing the megacohorts, see Methods) to the scaling factor estimated using synonymous mutations observed in all autosomal genes (Dig's default method), **a**, and using synonymous mutations observed in genes in the MSK IMPACT 230 targeted sequencing panel, **b**. **c,** The rate of excess missense SNVs with CADD phred<15 in tumor suppressor genes in the MSK IMPACT 230 targeted sequencing panel. The burden of missense SNVs with CADD phred<15 is not significant in any cancer type. **d,** The rate of excess pLoF SNVs in oncogenes in the MSK IMPACT 230 targeted sequencing panel. The burden of pLoF SNVs is not significant in any cancer type.

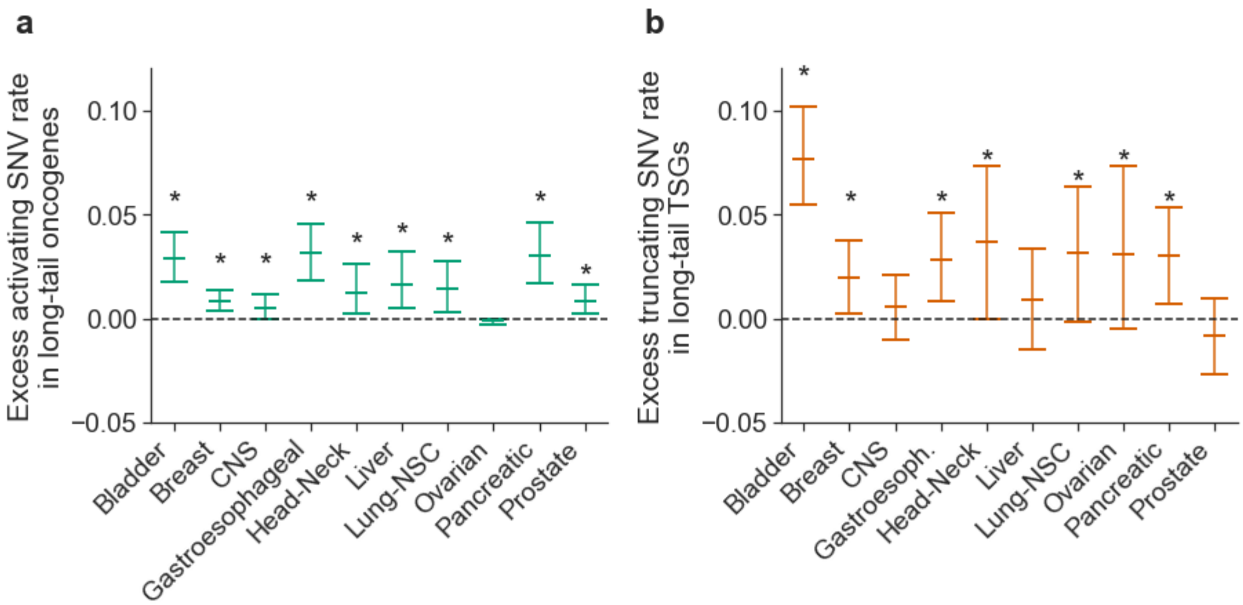

**Supplementary figure 18:** Excess activating SNVs in oncogenes, **a**, and excess pLoF SNVs in TSGs, **b**, as in **Fig. 4a,b** but with analysis restricted to whole-exome sequenced samples only. Asterisks indicate the burden of SNVs is significant in the given cancer type. Error bars are larger than in **Fig. 4a,b** because sample size is smaller (see **table S13** and **table S23** for exact sample sizes).

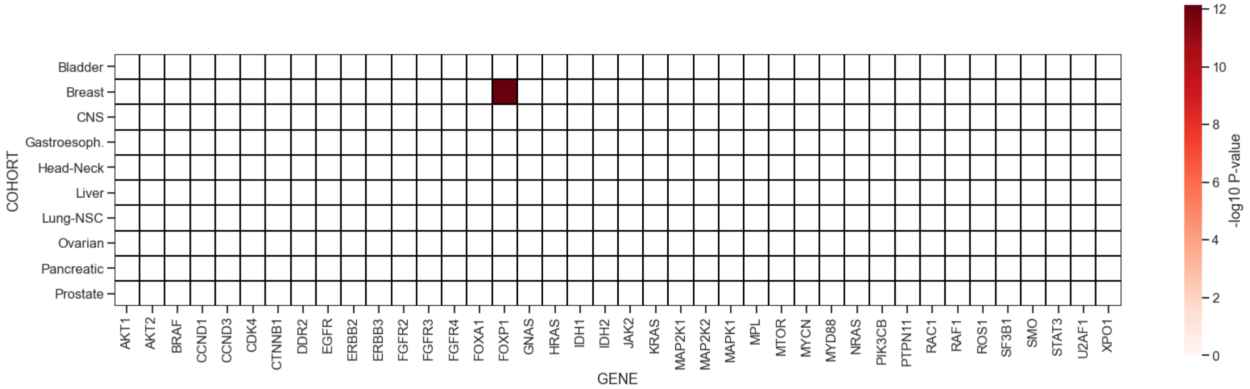

**Supplementary figure 19:** Burden of pLoF mutations in long-tail oncogenes. Oncogenes are not expected to have a significant burden of pLoF mutations.

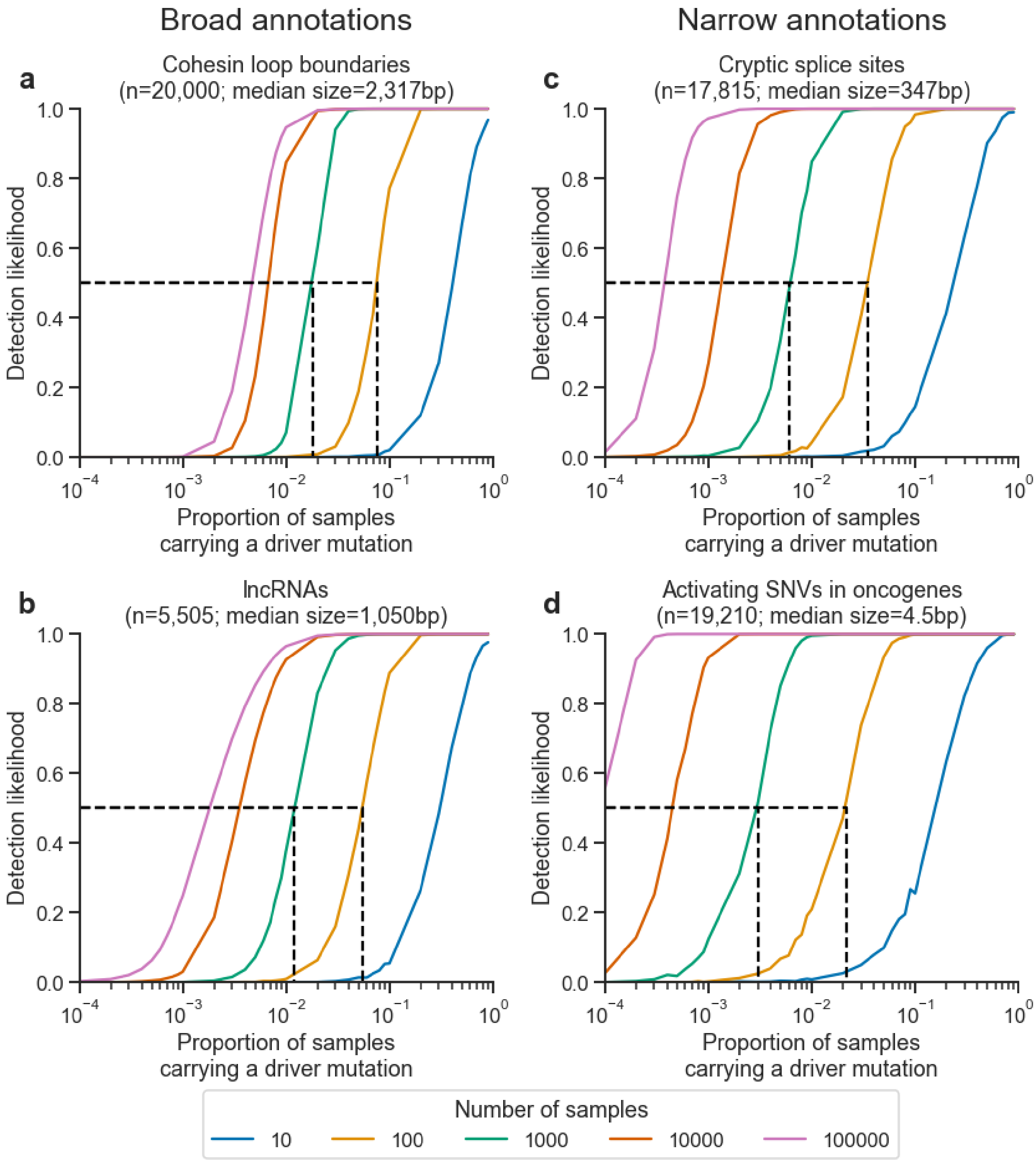

**Supplementary figure 20**: Driver detection likelihood based on the proportion of samples carrying a driver SNV for annotations containing thousands of base-pairs per element (broad annotations): cohesion loop boundary domains, **a**, and long noncoding RNAs (lncRNAs), **b**, and annotations containing tens to hundreds of base-pairs per element (narrow annotations): predicted cryptic intronic splice sites, **c**, and known activating SNVs in oncogenes, **d**). Dashed black lines indicated the proportion of samples required to carry a driver SNV to have a 50% likelihood of detecting the driver element in cohorts of 100 samples and 1000 samples, the approximate sample sizes of cancer cohorts used in this study.

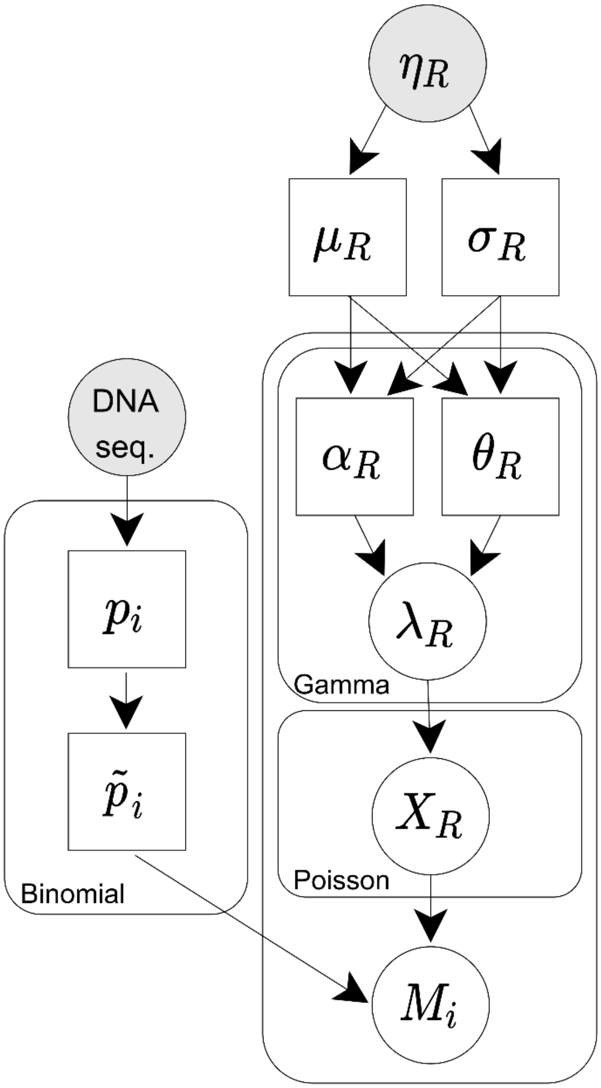

**Supplementary figure 21: Plate diagram of the probabilistic model that Dig uses to model the number of neutral mutations ($M_i$) in an element of interest**. $\eta_R$: observed data used as input to Dig's deep-learning model (chromatic modifications and, optionally, flanking mutation counts) to estimate regional neutral mutation parameters for region $R$. $\mu_R$ and $\sigma_R$: mean and standard deviation estimates of the neutral mutation rate in region $R$. $\alpha_R$ and $\theta_R$ gamma distribution shape and scale parameters, respectively. $\lambda_R$ gamma-distributed mutation rate parameter for region $R$. $X_R$ poisson-distributed mutation count in region $R$. DNA sequence (seq.): the DNA sequence from the human reference genome. $p_i$: genome-wide likelihood of a mutation in a given DNA context centered at position $i$. $\tilde{p}_i$: likelihood of mutation based on sequence context centered at position $i$ normalized such that $\sum_{i \in R} \tilde{p}_i = 1$. See **Methods** for additional details.

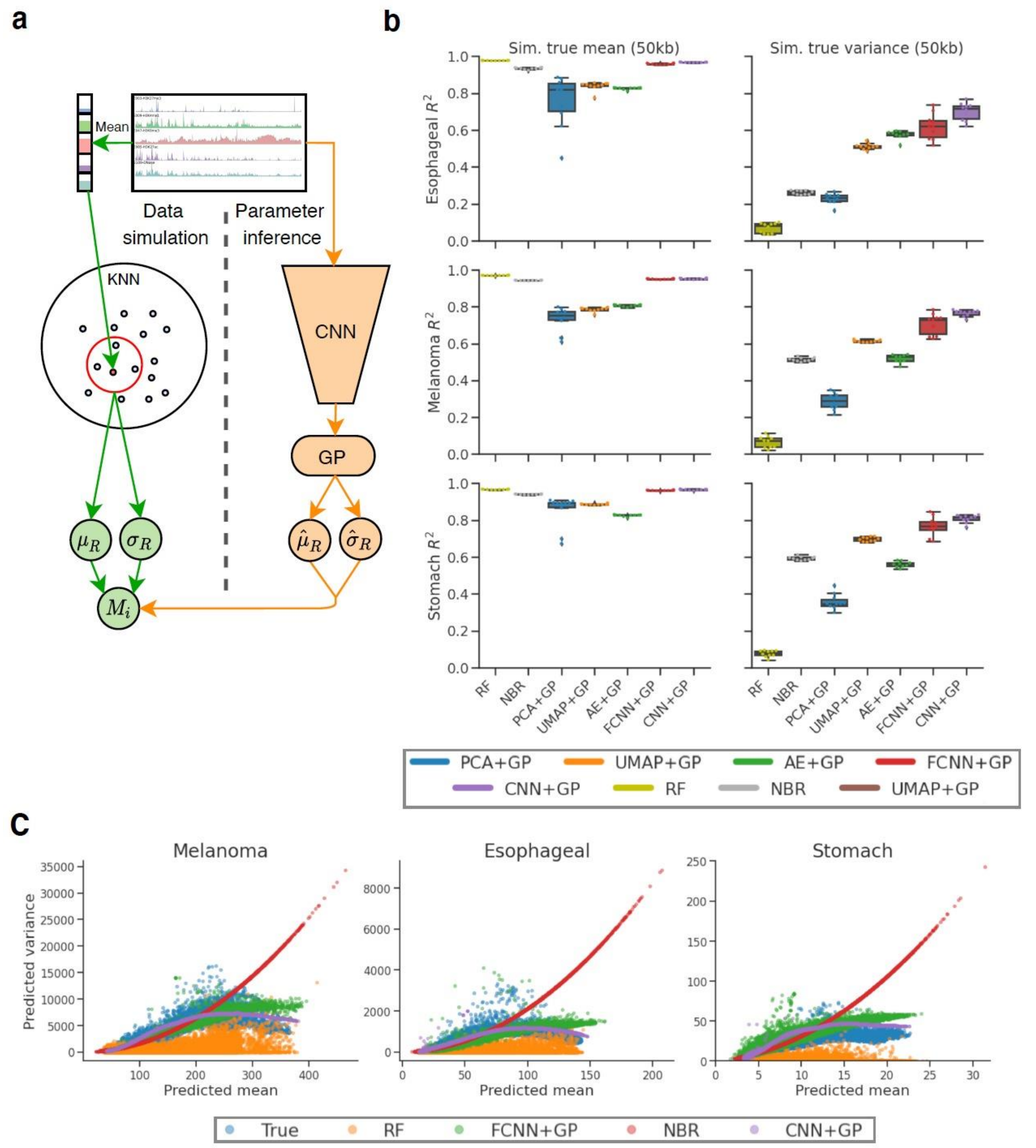

**Supplementary figure 22: Comparison of mean and variance prediction accuracy by method in simulated datasets. a,** Schematic of simulation framework. Briefly, for each 50kb region of the genome, a "true" mean and variance are constructed using a K-nearest-neighbors algorithm. The number of observed neutral mutations in that region is then simulated from a negative binomial distribution parameterized by this mean and variance. A method is then trained to predict the unknown mean and variance using the simulated number of mutations as a noisy objective. The accuracy of the model is evaluated by comparing the predicted mean and variance parameters to the known simulated values. **b,** Accuracy (Pearson's $R^2$) of the predicted mean and variance compared to the true simulated mean and variance across methods. RF: random forest; NBR: negative binomial regression; PCA+GP: principle components dimensionality reduction following by a Gaussian process; UMAP+GP: UMAP dimensionality reduction followed by a Gaussian process; AE+GP: autoencoder dimensionality reduction followed by a Gaussian process; FCNN+GP: fully connected neural network followed by a Gaussian process; CNN+GP: Convolutional neural network followed by a Gaussian process (Dig's default model). **c,** Mean versus variance of the simulated data (blue) and predicted by a CNN+GP (purple), FCNN+GP (green), negative binomial regression (red), or a random forest (orange).

14. Titsias, M. K. Variational Learning of Inducing Variables in Sparse Gaussian Processes. *AISTATS* 8 (2009).
15. Gardner, J. R., Pleiss, G., Bindel, D., Weinberger, K. Q. & Wilson, A. G. GPyTorch: Blackbox Matrix-Matrix Gaussian Process Inference with GPU Acceleration. *arXiv:1809.11165 [cs, stat]* (2019).
16. Supek, F. & Lehner, B. Scales and mechanisms of somatic mutation rate variation across the human genome. *DNA Repair (Amst)* **81**, 102647 (2019).
17. Elliott, K. & Larsson, E. Non-coding driver mutations in human cancer. *Nat Rev Cancer* 1–10 (2021) doi:10.1038/s41568-021-00371-z.
18. Yaari, A. U. *et al.* Multi-resolution modeling of a discrete stochastic process identifies causes of cancer. *International Conference on Learning Representations* (2020).
19. Shuai, S., PCAWG Drivers and Functional Interpretation Working Group, Gallinger, S., Stein, L., & PCAWG Consortium. Combined burden and functional impact tests for cancer driver discovery using DriverPower. *Nat Commun* **11**, 734 (2020).
20. Lochovsky, L., Zhang, J., Fu, Y., Khurana, E. & Gerstein, M. LARVA: an integrative framework for large-scale analysis of recurrent variants in noncoding annotations. *Nucleic Acids Res* **43**, 8123–8134 (2015).
21. Tate, J. G. *et al.* COSMIC: the Catalogue Of Somatic Mutations In Cancer. *Nucleic Acids Research* **47**, D941–D947 (2019).
22. Zhu, H. *et al.* Candidate Cancer Driver Mutations in Distal Regulatory Elements and Long-Range Chromatin Interaction Networks. *Molecular Cell* **77**, 1307-1321.e10 (2020).
23. Simonyan, K., Vedaldi, A. & Zisserman, A. Deep Inside Convolutional Networks: Visualising Image Classification Models and Saliency Maps. *arXiv:1312.6034 [cs]* (2014).
24. meuleman. *meuleman/epilogos*. (2021).
25. Kerpedjiev, P. *et al.* HiGlass: web-based visual exploration and analysis of genome interaction maps. *Genome Biology* **19**, 125 (2018).
